# Cortical astrocytes extend phasic norepinephrine signals and critically mediate learned behavior

**DOI:** 10.1101/2024.10.24.620009

**Authors:** Gabrielle T. Drummond, Arundhati Natesan, Marco Celotto, Jiho Park, Jennifer Shih, Prachi Ojha, Yuma Osako, Giselle Fernandes, Kyle R. Jenks, Grayson O. Sipe, Vincent Breton-Provencher, Tatsuya Osaki, Paul C. Simpson, Stefano Panzeri, Mriganka Sur

**Affiliations:** Department of Brain and Cognitive Sciences, Massachusetts Institute of Technology, Cambridge, MA, 02139, USA; Picower Institute for Learning and Memory, Massachusetts Institute of Technology, Cambridge, MA, 02139, USA; Institute for Neural Information Processing, Center for Molecular Neurobiology (ZMNH), University Medical Center Hamburg-Eppendorf (UKE), 20251 Hamburg, Germany; Department of Biology, Eberly College of Science and Huck Institutes of the Life Sciences, Pennsylvania State University, University Park, PA 16802, USA; Department of Psychiatry and Neuroscience, CERVO Brain Research Center, Université Laval, Québec City, Québec, Canada; Department of Medicine and Research Service, San Francisco Veterans Affairs Medical Center and Cardiovascular Research Institute, University of California, San Francisco, CA 94143, USA

## Abstract

**Updating behavior based on feedback from the environment is a crucial means by which organisms learn and develop optimal behavioral strategies**^1–3^**. Norepinephrine (NE) release from the locus coeruleus (LC) has been shown to mediate learned behaviors**^4–6^ **such that in a task with graded stimulus uncertainty and performance, a high level of NE released after an unexpected outcome causes adaptations in subsequent behavior**^7^**. Yet, how the transient activity of LC-NE neurons, lasting tens of milliseconds, alters neuronal activity and influences behavior several seconds later is unclear. Here, we show that NE released after an unexpected outcome acts directly on cortical astrocytes via** α1 **adrenergic (Adra1a) receptors to elicit sustained increases in intracellular calcium. Chemogenetic blockade of astrocytic calcium dynamics prevents trial-to-trial behavioral adaptation. NE stimulation of astrocytes elicits ATP release, and imaging ATP levels in the cortex reveals an increase in extracellular ATP in response to an unexpected outcome. Blocking ATP-driven signaling to neuronal adenosine A1 receptors also prevents post-reinforcement behavioral adaptation. Finally, high density neuronal recordings in prefrontal cortex reveal that a surprising outcome alters the neuronal representation of the stimulus on the subsequent trial without sustained changes in cortical activity; blocking either astrocyte calcium dynamics or A1 receptors occludes these post-reinforcement changes in single-neuron and population neuronal encoding of task variables underlying behavioral changes. Together, these data demonstrate that astrocytes play an essential role in norepinephrine-driven learned behavior: they have prolonged calcium responses to transient norepinephrine release and convey task-relevant reinforcement information across behavioral intervals, enabling selective updating of neuronal task representations to support adaptive behavior.**

## Introduction

Norepinephrine (NE) released by locus coeruleus (LC) neurons is associated with diverse functions, including regulation of arousal and sleep^8–11^, stress^12–14^, memory consolidation, and learning^7,15–18^. These varying functions of LC-NE are thought to be enabled by different modes of LC activity^11,15,19,20^, a semi-modular organization with diverse inputs and targets of LC-NE projections^7,15,21–26^, and by behavior dependent processing in target regions based on receptor expression. While slowly varying activity of LC-NE neurons has been linked to locomotion and arousal, task-dependent transient or phasic LC-NE activity acts as a selective learning signal, complementary to the non-selective role of NE in arousal^7,15,23,27–34^. A crucial function of phasic LC-NE activity during reinforcement-based learned behavior is to signal prediction error in order to optimize behavior^6,7,15,32,35,36^. Yet, how a brief LC-NE prediction error signal reorganizes cortical responses to alter behavior on longer time scales is unknown.

Astrocytes are the major non-neuronal cell type in the cortex and are increasingly recognized as key contributors to the development, plasticity, and function of neuronal circuits^37–44^. They are highly responsive to NE^45–50^ and exhibit calcium dynamics ranging from hundreds of milliseconds to several seconds, suggesting that these signals can reflect as well as influence neuronal activity and behavior on a range of time scales. Yet, whether and how astrocytes might dynamically shape cortical circuits underlying learning and learned behavior is poorly understood. Here, we show that in a learned behavioral task, cortical astrocytes in prefrontal and motor cortex are highly responsive to phasic NE released after a reinforcement, and their prolonged calcium dynamics causally drive behavioral updating on the next trial. NE-driven astrocyte modulation of behavior involves purinergic signaling to neurons, which alters neuronal stimulus encoding on trials following a surprising outcome and consequent negative or positive reinforcement. Thus, astrocytes have an essential role in NE-dependent learned behavior: they integrate NE reinforcement signals and convey them across behavioral intervals to modulate neuronal activity and mediate behavioral adaptation.

## Results

### Astrocytes exhibit sustained increases in calcium following an unexpected outcome

To evaluate how phasic NE reinforcement signals affect cortical processing to mediate behavioral adaptation, we trained mice in a go/no-go task with graded auditory stimulus evidence where animals must press a lever at a go tone to receive a water reward and refrain from pressing at a no-go tone to avoid an air puff punishment^7^ (**Figures 1A, S1A, and S1B**). We have shown previously that LC-NE neurons exhibit phasic activity during two distinct task epochs, pre-press and post-reinforcement^7^ (**Figure 1B**). Using 2-photon imaging of an NE sensor, we confirmed that NE is released in the cortex at these two epochs for both hit and false alarm trials (**Figures S1C-S1F**). This LC-NE activity plays dual roles in the task: low phasic LC-NE activity pre-press facilitates task execution on low stimulus evidence trials, while high phasic LC-NE activity following an unexpected outcome, such as a false alarm followed by an air puff or a correct rejection followed by a reward, drives systematic changes in behavior on the subsequent trial, several seconds later^7^ (**Figures 1B,1C, and 1D**). On hit trials, pre-press LC-NE neuronal activity correlates with tone intensity, while post-reward activity negatively correlates with go tone intensity, consistent with prediction error encoding^7^. Accordingly, after a false alarm, the most unexpected outcome, LC-NE neurons exhibit their highest activity. Indeed, on trials after this high phasic NE activity, the stimulus-response psychometric curve is shifted such that mice perform fewer false alarms and more hits on the next trial, resulting in a serial response bias which improves response sensitivity or d-prime, the ability to discriminate between go and no-go stimuli, on the subsequent trial (**Figures 1C and 1D**). The increase in d-prime following a false alarm is also significantly higher than after a hit or unreinforced (miss and correct rejection) trial (**Figure S1G**). Thus, a phasic LC-NE signal following a surprising outcome encodes reinforcement surprise to facilitate behavioral performance on the next trial, several seconds later.

**Figure 1.**
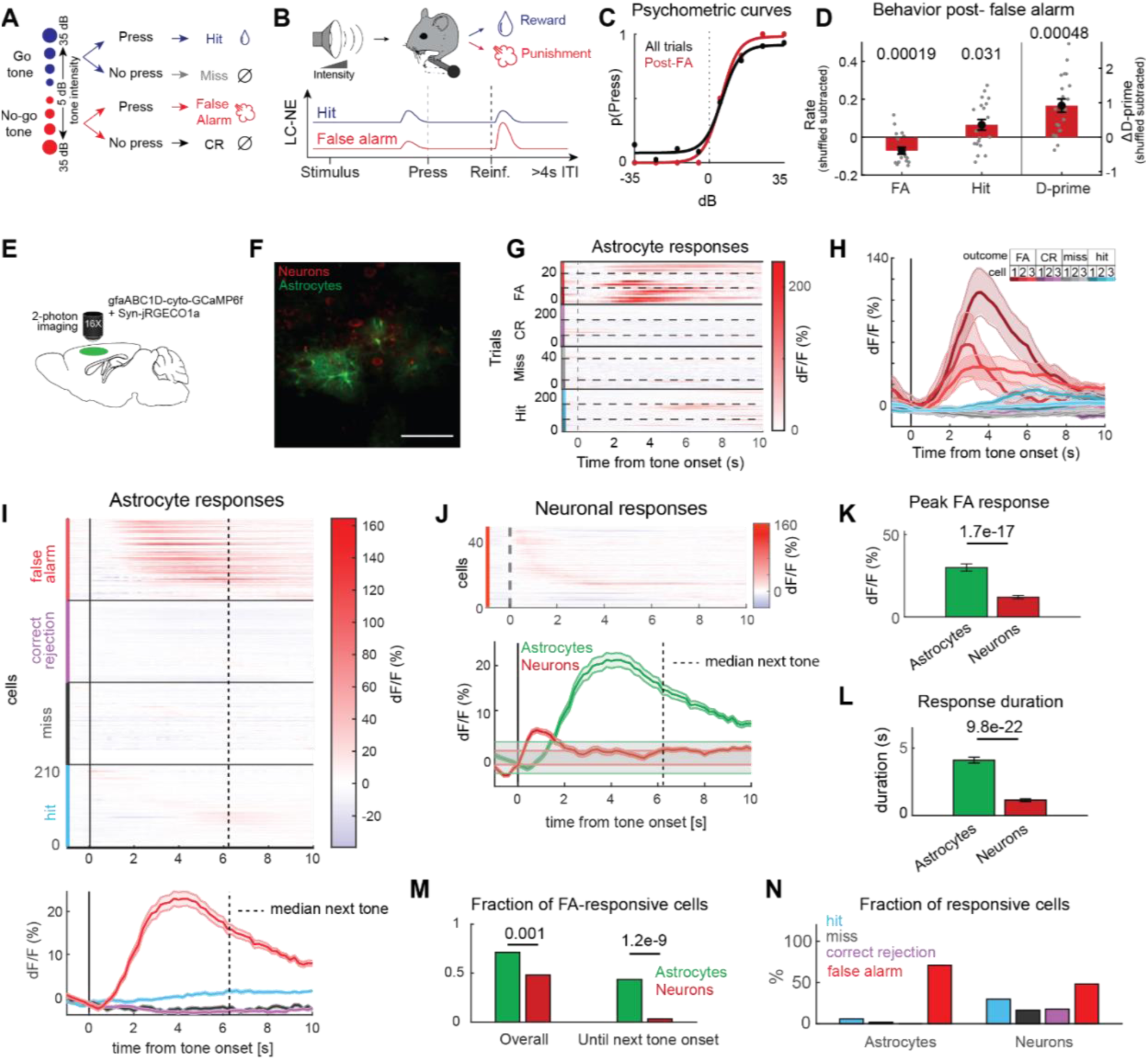
Astrocytes exhibit sustained increases in intracellular calcium following a false alarm in a reinforcement learning task. (**A**) Task design. **(B)** Schematic of task and timing/magnitude of LC-NE neuronal activity. ITI, inter-trial interval. **(C)** Example psychometric curves from one mouse showing probability of pressing the lever by tone intensity for all trials (black) and on trials following a false alarm (red). **(D)** Change in false alarm rate, hit rate, and d-prime on trials following a false alarm, calculated after subtracting shuffled data (n = 20 mice). **(E)** Methods for dual imaging of astrocytes and neurons. **(F)** Example two-photon image of astrocytes and neurons expressing GCaMP6f and jRGECO1a, respectively. Scale bar = 50 µm. **(G)** Mean dF/F of three example astrocytes aligned to tone onset for false alarm (FA), correct rejection (CR), miss, and hit trials. **(H)** Session averaged dF/F for the three example astrocytes aligned to tone onset, by trial type. **(I)** Top: raster plot showing average astrocyte dF/F by trial type, aligned to the time of tone inset. Bottom: average dF/F for all astrocyte ROIs by trial type. N = 218 astrocytes (n = 4 mice). **(J)** Top: raster plot showing average neuronal dF/F on false alarm trials. Bottom: Population average dF/F for all neurons and all astrocytes (from **i**) on false alarm trials, aligned to time of tone onset. Gray zones depict baseline ± s.d. n = 208 neurons (n = 3 mice). **(K)** Peak dF/F for astrocytes and neurons in response to a false alarm. **(L)** Duration of astrocyte and neuron responses to a false alarm. **(M)** Fraction of astrocytes and neurons responsive to a false alarm (left) and whose responses lasted until the next median tone onset (right). **(N)** Fraction of astrocytes (left) and neurons (right) responsive to each trial type. P values show comparisons based on two-tailed one-sample t-test in **D**, two-tailed unpaired t-test in **K**, **L**, and two-tailed normal approximation to binomial test in **M**. Data show mean ± SEM. Gray and black vertical dashed lines in **I**, **J** show the median time of next tone onset.

Due to their spatiotemporally diverse activity patterns and their proximity to tens of thousands of synapses per astrocyte^51,52^, we hypothesized that astrocytes could integrate phasic reinforcement signals with local neuronal activity to alter neuronal population dynamics and improve performance after such a prediction error. Notably, astrocytes express α1 adrenergic (Adra1a) receptors and have been shown to be responsive to NE release during vigilance and changes in internal states^53^. Astrocytes have been suggested to mediate switches in cortical states in response to NE signaling^54^, but whether and how NE signaling plays a role in learned behaviors remains unknown. To examine astrocyte activity in response to controlled NE release *in vivo*, we first used two-photon imaging of astrocyte calcium after optogenetic activation of channelrhodopsin (ChR2)-expressing NE axons through a cranial window in the visual cortex (**Figures S2A and S2B**). Astrocyte calcium signals increased sharply with NE release in the cortex, such that longer stimulation paradigms elicited significantly larger increases in calcium responses (**Figures S2C and S2D**). Thus, cortical astrocytes are indeed responsive to NE release *in vivo*, with higher calcium elevations and potentially more astrocytes recruited in response to higher NE release.

To examine the effects of NE on astrocytes and neurons in target regions during behavior, we used dual two-photon imaging of astrocytic and neuronal calcium responses in the motor cortex of mice performing the go/no-go task (**Figures 1E and 1F**). We have previously shown that prefrontal cortex and motor cortex are involved in the task and receive similar NE signals following a false alarm^7^. We found that, on average, motor cortex astrocytes exhibited consistent increases in calcium activity in reinforced trials compared to unreinforced trials with a larger increase following a false alarm air puff (**Figures 1G-1I and S2E-S2G**), as well as following a reward after a correct rejection (see below), both being unexpected task outcomes that lead to phasic NE release. Notably, we observed that astrocytes strongly responsive to false alarm trials also showed greater sensitivity to tone intensities in hit trials, with post-hit responses exhibiting a negative correlation with go tone intensity, (**Figures S2H and S2I**), a hallmark of reward prediction error signaling that recapitulated similar responses of LC-NE neurons^7^. Thus, a wide range of reinforcement-driven phasic LC-NE responses lead to astrocyte calcium signaling. We observed heterogeneity in astrocyte calcium responses to a false alarm (**Figure 1H**), with some astrocytes exhibiting higher peak elevations in calcium than others (**Figure S2E**). Astrocyte responses correlated with the probability of making the correct choice on the subsequent trial (**Figure S2J**), suggesting that astrocytes causally influence task performance.

Comparing astrocyte calcium increases with neuronal calcium increases during the task, we found that neuronal responses had faster rise times but shorter durations, shorter than the inter-trial interval (>4 s), in our task (**Figures 1J, S1A, and S1B**), suggesting that persistent neuronal activity alone is not sufficient to bridge between one trial and the next. Astrocyte calcium responses were significantly larger and remained elevated for significantly longer durations, spanning the inter-trial interval (**Figures 1J-1M**). Finally, while neurons exhibited heterogeneous activity during the task and were responsive to multiple trial types, astrocytes were strongly responsive to false alarm trials and also exhibited responses to hit trials, albeit of smaller magnitude (**Figures 1N and S2E-S2I**). We performed clustering analysis of astrocyte calcium responses in reinforced trial types and characterized two astrocyte populations, the larger of which displayed strong, sustained responses to false alarms (**Figures S2K-S2M**). These data demonstrate that astrocytes are highly responsive to a surprising air puff reinforcement following a false alarm, and have long-lasting calcium increases which correlate with next trial performance, unlike cortical neurons which exhibit shorter responses that do not correlate with next trial performance (**Table S1**).

To examine whether astrocyte calcium increases might be due to the aversive nature of the air puff, we measured astrocyte responses to a surprising reward. Giving a water reward on a correct rejection trial, which is typically unreinforced, has been shown to elicit high LC-NE activity similar to that following a surprising air puff on a false alarm trial^7^. Two-photon imaging of the NE sensor while a surprising reward was given on a subset of correct rejection trials revealed that extracellular cortical NE release was as high following a rewarded correct rejection as following a false alarm air puff (**Figures S3A-S3D**). We then measured astrocyte calcium dynamics during the task (**Figure S3E**) and performed clustering analysis of their responses in the trials with unexpected reinforcements: we observed two astrocyte populations, both with large response magnitudes but exhibiting a stronger response for one trial type over the other, suggesting functional heterogeneity in how astrocytes encode different types of surprising events (**Figures S3F and S3G**). These data demonstrate that surprising outcomes for both positive and negative reinforcement that cause high NE release can also strongly activate astrocytes.

Since NE activity has been associated with changes in arousal states^54,55^, as characterized by an increase in high frequency power in neuronal local field potential (LFP) activity in the cortex^54^ and by changes in pupil diameter^56,57^, we next evaluated these measures to determine if changes in behavior following a false alarm could be explained by arousal alone. We analyzed LFP activity in the cortex and measured the power across all frequencies including low frequency (2-30 Hz) and gamma frequency (30-80 Hz) bands. We observed low frequency and gamma power responses following a false alarm as well as after a hit, but these responses were not sustained across the inter-trial interval^54,58^ (**Figures S4A-S4C**), indicating that false alarm-induced changes in such activity are short-lasting and are unlikely to mediate persistent brain state changes that could underlie behavioral gain on the next trial. Furthermore, average low or gamma frequency power did not correlate with the probability of a correct response on the next trial after a false alarm (**Table S1**), suggesting that LFP dynamics do not mediate behavioral adaptation in this task. In addition, we observed no significant relationship between post-false alarm pupil diameter change and the probability of success on the next trial (**Table S1**). Taken together, these data indicate that surprising outcomes elicit transient changes in cortical neuronal activity that do not span the inter-trial interval nor correlate with next trial performance and thus are unlikely to account for NE-driven behavioral optimization. This is in line with our findings above and previous data supporting a specific role for NE in mediating reinforcement learning through phasic NE release to signal uncertainty and prediction errors, separable from its role in mediating arousal^4,7,15,59–61^.

### NE-astrocyte signaling mediates improvements in behavioral performance

To investigate whether astrocyte calcium increases following a false alarm were dependent on NE release, we measured astrocyte calcium responses in mice while reversibly chemogenetically silencing LC-NE neurons. We expressed a floxed Gi-coupled designer receptor exclusively activated by designer drugs (DREADD) in Dbh-Cre mice and imaged astrocyte calcium 30 minutes after injecting low-dose CNO or saline (**Figures 2A and 2B**). When Gi-DREADDs in the LC were activated using CNO, astrocytes in the motor cortex were less responsive to a false alarm than during saline sessions, exhibiting lower magnitude peak and average responses and shorter duration of responses, suggesting that astrocyte calcium increases reflect LC-NE signaling (**Figures 2C, 2D, and S5A-S5C**). Because optogenetic silencing of LC-NE neurons prevents the behavioral gain of false alarm punishments on the next trial^7^ and astrocyte false alarm responses correlate with next trial behavioral performance, we hypothesized that astrocyte calcium increases in response to phasic LC-NE release are causal for the behavioral gain.

**Figure 2.**
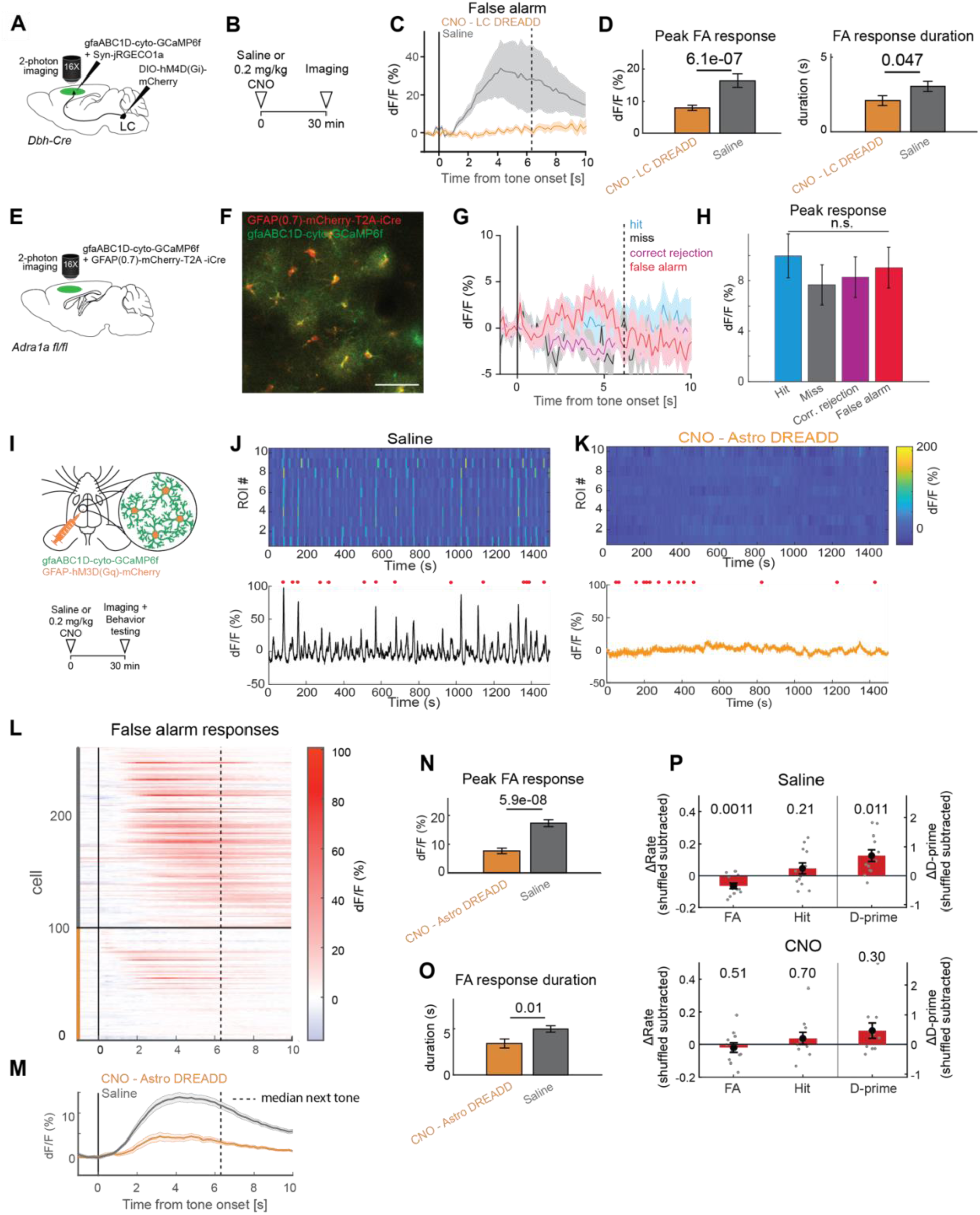
NE and adrenergic receptor-specific responses of astrocytes and effects of manipulating calcium on behavior. **(A)** Experimental design for chemogenetic silencing of LC-NE neurons while imaging astrocyte calcium. **(B)** Experimental timeline. **(C)** Example trial averaged astrocyte trace on a false alarm trial for a saline and CNO session. **(D)** Left: Comparison of peak astrocyte dF/F on false alarm trials for saline and CNO sessions. Right: Average false alarm response duration for responsive astrocytes in saline and CNO sessions. n = 108 astrocytes, CNO; n = 101 astrocytes, saline. (n = 3 mice). P values show comparisons based on two-tailed unpaired t-test. **(E)** Experimental design for imaging astrocyte calcium in astrocytes with Adra1a receptor knockdown (KD). **(F)** Example two-photon image showing overlapping Cre and GCaMP6f expression in astrocytes. Scale bar = 50 µm. **(G)** Example traces for hit, miss, correct rejection, and false alarm trials from one astrocyte. **(H)** Population averages for peak astrocyte dF/F for all trial types in Adra1a-R KD astrocytes. n = 44 astrocytes. (n = 3 mice). **(I)** Experimental design and timeline for chemogenetic manipulation of astrocyte calcium. **(J)** Top: Average dF/F for each astrocyte ROI over the course of a saline imaging session. Bottom: population average. Red dots indicate false alarms. **(K)** Average dF/F for each astrocyte ROI over the course of a CNO imaging session. Bottom: population average. Red dots indicate false alarms. **(L)** Average astrocyte dF/F on false alarm trials for saline (top) and CNO (bottom) sessions. n = 99 astrocytes, CNO; n = 160 astrocytes, saline (n = 5 mice). **(M)** Population average from **l**. **(N)** Average peak dF/F for astrocytes in saline and CNO sessions. **(O)** Average response duration for responsive astrocytes in saline and CNO sessions. **(P)** Change in false alarms (FA), hits, and d-prime on trials after a false alarm for saline (top) and CNO (bottom) sessions. (n = 12 mice). P values show comparisons based on two-tailed unpaired t-test in **D, N, O,** one-way ANOVA in **H** (p = 0.054), and two-tailed one-sample t-test in **P**. Data show mean ± SEM. Gray and black vertical dashed lines in **C**, **G**, **L**, **M** show the median time of next tone onset.

Since astrocytes could also be responding to activity in nearby neurons following a false alarm, we next evaluated whether NE binding to adrenergic receptors on astrocytes is critical for the increase in calcium following a false alarm. Using *Adra1a fl/fl* mice, we expressed GFAP(0.7)-mCherry-T2A-iCre and gfaABC1D-GCaMP6f in cortical astrocytes to selectively knock down Adra1a receptors in a subset of cortical astrocytes and imaged the calcium activity in the same astrocytes during the task (**Figures 2E and 2F**). To confirm effective knockdown of Adra1a expression, we used fluorescence *in situ* hybridization (FISH) to label *Adra1a*, *Slc1a3* (mRNA for the astrocyte specific glutamate transporter, EAAT1/Glast1), and *mCherry* mRNA in the prefrontal and motor cortex (**Figure S6A**). We found heterogenous *Adra1a* mRNA expression in astrocytes which was nevertheless significantly higher than in non-astrocytic cells (**Figure S6B**). The varying expression of Adra1a in astrocytes may partially explain the heterogeneity of astrocytic calcium responses to optogenetic activation of ChR2-expressing NE axons (**Figures S2C and S2D**) and to surprising outcomes in the task (**Figures S2E-S2G, S2K-S2M, and 3F)**. In comparing *mCherry* positive and negative astrocytes from the same animals, we saw a significant reduction in the number of detected *Adra1a* puncta in *mCherry* positive cells, confirming effective knockdown (**Figure S6C**). Without Adra1a receptors, cortical astrocytes no longer exhibited increases in calcium following a false alarm, with calcium response magnitudes similar to other trial outcomes (**Figures 2G and 2H**). We compared astrocyte calcium responses in *Adra1a fl/fl* mice to mice with no manipulations and saw a significant decrease in false alarm response magnitude in the Adra1a receptor knockdown mice (**Figures S7A-S7E**). These data indicate that astrocytes respond directly to NE release post-false alarm via Adra1a receptors and not indirectly by reflecting the activity of nearby neurons.

To establish whether activation of Adra1a receptors on astrocytes is important for learning the task, we next evaluated whether these mice exhibited behavioral deficits during training. After completing stage one of training (associating go tones with lever press and water reward), mice with Adra1a receptor knockdown in the PFC and MC exhibited similar hit rates as wild-type animals when they progressed to stage two (introduction of no-go tones; see Methods) (**Figures S7F and S7G).** However, while wild-type mice did fewer false alarms as training progressed in stage two, suggesting that they started to associate pushing the lever on no-go tones with receiving an air-puff, Adra1a receptor knockdown mice continued to have a high false alarm rate (**Figures S7F and S7G**). Thus, while wild-type mice showed a rapid day-by-day improvement in d-prime over the early days of training with no-go tones, the Adra1a knockdown mice did not, suggesting that Adra1a receptor activation on astrocytes is essential for a key component of task acquisition.

In addition to this chronic manipulation, we sought to use a session-based manipulation, using the same animals as controls, to ask whether the increase in astrocyte calcium following a false alarm is critical for the behavioral improvement observed after a false alarm. To deplete intracellular calcium stores and blunt astrocyte calcium transients following a false alarm, as shown previously^62,63^, we expressed Gq-DREADD in astrocytes in the prefrontal and motor cortex of wild-type mice trained in the task and activated the DREADDs using i.p. injections of low-dose CNO 30 minutes prior to behavioral testing and imaging (**Figure 2I**). Consistent with our previous findings^63^, when we injected saline as control we saw dynamic astrocyte calcium activity over the course of the session, with large peaks in population activity largely correlating with false alarms (**Figure 2J**). When we activated the DREADD however, astrocyte calcium activity no longer showed dynamic changes (**Figure 2K**). Indeed, when we aligned calcium responses to the behavior, astrocyte calcium activity following a false alarm was highly attenuated (**Figures 2L and 2M**) and showed a significant reduction in the response amplitude and duration compared with saline controls (**Figures 2N and 2O**). We observed variations in astrocyte calcium responses following other trial types, but calcium response levels in CNO false alarm trials were consistently reduced (**Figures S8A-S8E and 2L-2O**). Thus, Gq-DREADD activation effectively disrupted astrocyte calcium dynamics, enabling us to test the necessity of astrocyte calcium transients for behavioral gain in the task. We next assessed the effects of astrocyte calcium manipulation via Gq-DREADD on serial response bias observed in the task. In saline sessions, mice performed fewer false alarms and had improved response sensitivity or d-prime on trials after a false alarm, similar to wild-type animals, while in CNO sessions in the same mice this improvement in performance following a false alarm was specifically prevented (**Figures 2P and S9A**). Importantly, we found little generalized change in behavior as measured by overall hit rate, false alarm rate, and d-prime, indicating lack of non-specific effects following CNO administration (**Figure S9B**). Taken together, these data suggest that specific temporal dynamics in prefrontal and motor cortex astrocyte calcium are necessary for the improvement in response sensitivity on post-false alarm trials.

We next evaluated the effects of astrocyte calcium manipulation on behavior following a surprising reward. Following an unexpected water reward after correctly withholding a lever press, animals reduced their pressing on the subsequent trial, a behavioral effect that was not observed in trials following unrewarded correct rejection trials or false alarm trials (**Figures S10A-S10K**). Blocking astrocyte calcium dynamics by activating Gq-DREADDs in astrocytes abolished this behavioral effect following a rewarded correct rejection (**Figures S10A-S10H**). Although false alarm air puffs and correct rejection water rewards evoked similarly elevated NE activity (**Figures S3A-S3D**), they produced distinct behavioral consequences on the subsequent trial: a reduction in hits and drop in press probability or response bias following rewarded correct rejections versus an increase in hits and d’ following false alarms. This dissociation between similar NE elevations and different behavioral outcomes suggests that astrocytes may reflect the context in which NE release occurs, contributing to subsequent behavioral adaptation.

### Astrocytes mediate behavioral gain via purinergic signaling to neurons

Several mechanisms have been proposed for how astrocyte calcium signals influence synaptic and neuronal responses. Astrocytes can modulate extracellular K+^40^ and drive purinergic signaling by release of ATP^64–71^ under diverse conditions including in response to NE^72–75^. ATP is converted to adenosine extracellularly^76,77^ where it binds to adenosine-selective A1 receptors on neurons to modulate synaptic transmission^67,78–81^. Indeed, their robust response to NE and consequent influence on synaptic signaling, while examined only in slices^64,74^, suggests a mechanistic role for astrocytes in NE-dependent learning and behavior, but such a role has yet to be demonstrated causally^82^. We thus hypothesized that in our task, cortical astrocytes that are activated by NE following a false alarm could influence neuronal activity on the next trial via purinergic signaling^64–75,78–81^. To address this hypothesis, we first measured adenosine release in astrocyte cultures in response to NE using a fluorometric assay. Upon addition of NE, we saw a significant increase in adenosine fluorescence in the media. Blocking Adra1a receptors with the selective antagonist prazosin prevented the increase in adenosine fluorescence (**Figure 3A**), suggesting that NE signaling to astrocytes can cause increases in extracellular adenosine.

**Figure 3.**
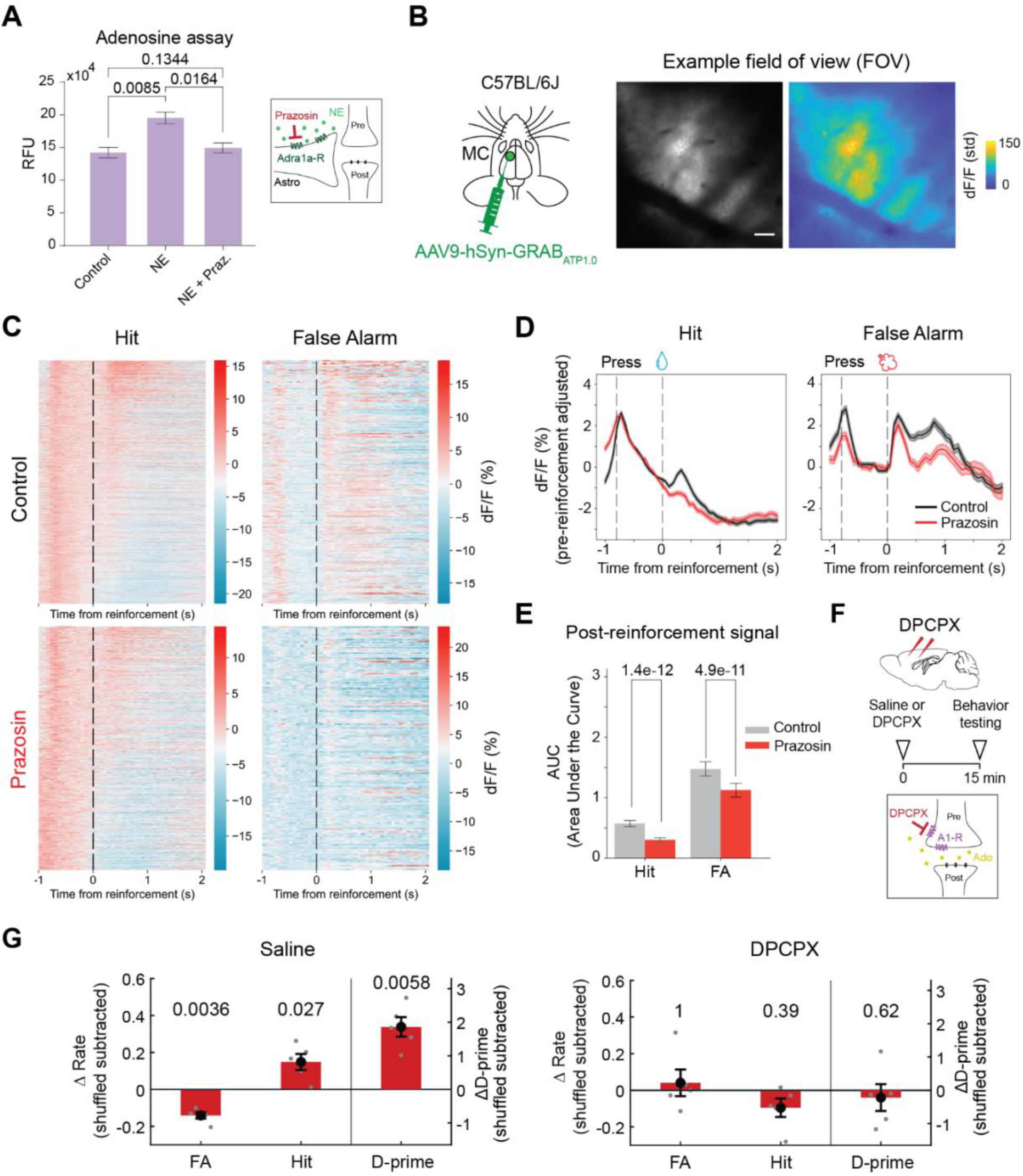
Astrocytes mediate reinforcement learning via purinergic signaling. **(A)** Adenosine fluorometric assay of astrocyte cultures showing an increase in adenosine in response to NE (20µM). NE-induced release of adenosine is blocked in the presence of prazosin hydrochloride (inset). RFU = relative fluorescence units. (n = 3 cultures). **(B)** (Left) Experimental design for *in vivo* imaging of GRAB_ATP_ sensor in motor cortex. (Right) Example field of view of two-photon imaging session. Right panel shows a pseudocolor image (STD = standard deviation). Scale bar = 50 µm. **(C)** Heatmaps of baseline-adjusted GRAB_ATP_ dF/F traces in hit and false alarm trials. Individual trial signals were aligned to the time of reinforcement (indicated in black dashed line). (n = 3 mice). **(D)** Mean dF/F trace averaged across all trials for each trial type. GRAB_ATP_ dynamics after systemic administration of prazosin hydrochloride are in red. **(E)** Quantification of post-reinforcement signals. AUC values were computed over a fixed time window after reinforcement delivery. Hit: Control = 1756 trials from 3 mice. Prazosin = 1004 trials from 3 mice. False Alarm: Control = 395 trials from 3 mice. Prazosin = 226 trials from 3 mice. **(F)** Experimental design for applying saline or DPCPX topically to prefrontal and motor cortex, to block A1 receptors (inset), followed by behavioral testing. **(G)** Change in false alarms, hits, and d-prime on trials after a false alarm in saline (left) and DPCPX (right) conditions. (n = 5 mice). P values show comparisons based on one-tailed unpaired t-test in **A**, **E**, and two-tailed one-sample t-test in **G**. Data show mean ± SEM.

We next asked whether NE release in the cortex during the behavioral task elicited increases in ATP *in vivo*. To evaluate the temporal dynamics of ATP release in the cortex, we used two-photon imaging of an ATP sensor in the motor cortex^83^ (**Figure 3B**). By aligning ATP measurements to the time of reinforcement, we found sharp, transient increases in ATP, both pre-press and post-reward, in hit trials, and a relatively broad ATP response following an air puff in false alarm trials (**Figures 3C and 3D**). Systemic administration of prazosin led to a significant reduction of post-reinforcement responses, particularly in the response amplitude and duration after false alarm trials (**Figures 3C-3E**). When we introduced a surprising reward following a correct rejection, we again observed an increase in ATP levels which was also significantly decreased upon prazosin administration (**Figures S11A-S11C**). These data indicate that NE action on astrocytes can elicit astrocyte release of ATP in response to a surprising outcome.

We therefore asked whether the action of ATP/adenosine on neurons mediates behavioral improvement in our task by assessing the effects of blocking adenosine A1 receptors in motor and prefrontal cortex *in vivo* in mice performing the task. When we applied DPCPX, a selective A1 receptor antagonist, topically to these cortical regions, mice no longer exhibited an increase in d-prime after a false alarm (**Figures 3F, 3G and S12A**). This effect on behavior was response-specific, as we did not observe generalized change in behavior as measured by overall hit rate, false alarm rate, and d-prime across conditions (**Figure S12B**). Additionally, DPCPX application abolished the decrease in lever press bias observed after rewarded correct rejection trials, similar to astrocyte calcium manipulations (**Figures S13A-S13J**). Overall, our findings suggest that purinergic signaling from astrocytes to neurons is critical for the behavioral changes observed after an unexpected outcome in our task.

### Astrocyte calcium manipulations and A1 receptor blockers alter post-reinforcement neuronal encoding of task variables

Finally, we asked how neuronal computations in cortex were influenced by a false alarm and subsequent air puff (or correct rejection-reward), and how manipulating astrocyte calcium increase and blocking astrocyte-neuron purinergic signaling in response to NE affected these computations. To address these questions, we generated high density neuropixel recordings of neurons in the prefrontal cortex of mice performing the task. We recorded neurons from the same mice with either saline, CNO activation of Gq-DREADD in cortical astrocytes, or topical application of the A1 receptor antagonist DPCPX (**Figure 4A**). Using the same mice with enough “wash-out” time between experiments enabled direct comparison between manipulations. Importantly, there was no change in overall firing rate as compared across saline, CNO, or DPCPX conditions or in post-false alarm firing rate compared with a shuffled-trial-order control within each condition (**Figures 14A and 14B**), and no generalized change in behavior as measured by hit rate, false alarm rate, and d-prime across conditions (**Figures S9A, S9B, S12A, and S12B**). Using a linear support vector machine decoder to classify false alarm trials from other trial types from the neuronal activity, we found that the time-resolved decoder performance accuracy dropped to chance level by the next trial start time for all conditions and did not correlate with next trial performance (**Table S1**, **Figure S14C**). Furthermore, similar to our analysis of post-reinforcement LFP power across all frequencies in wild-type mice (**Figure S4**), we analyzed post-reinforcement LFP power in Gq-DREADD and DPCPX conditions (**Figure S15**), and found no difference in the LFP power in any frequency band between wild-type mice or saline controls and Gq-DREADD or DPCPX treated mice after false alarms or rewarded correct rejection trials. Taken together, these analyses indicate that several components of post-reinforcement neuronal activity (single-cell firing rate levels, LFP power, and population false alarm signals) are unaffected by Gq-DREADD or DPCPX manipulations, which we have shown to block changes in performance following a surprising outcome, suggesting that these neuronal signals are unlikely to mediate the behavioral changes.

**Figure 4.**
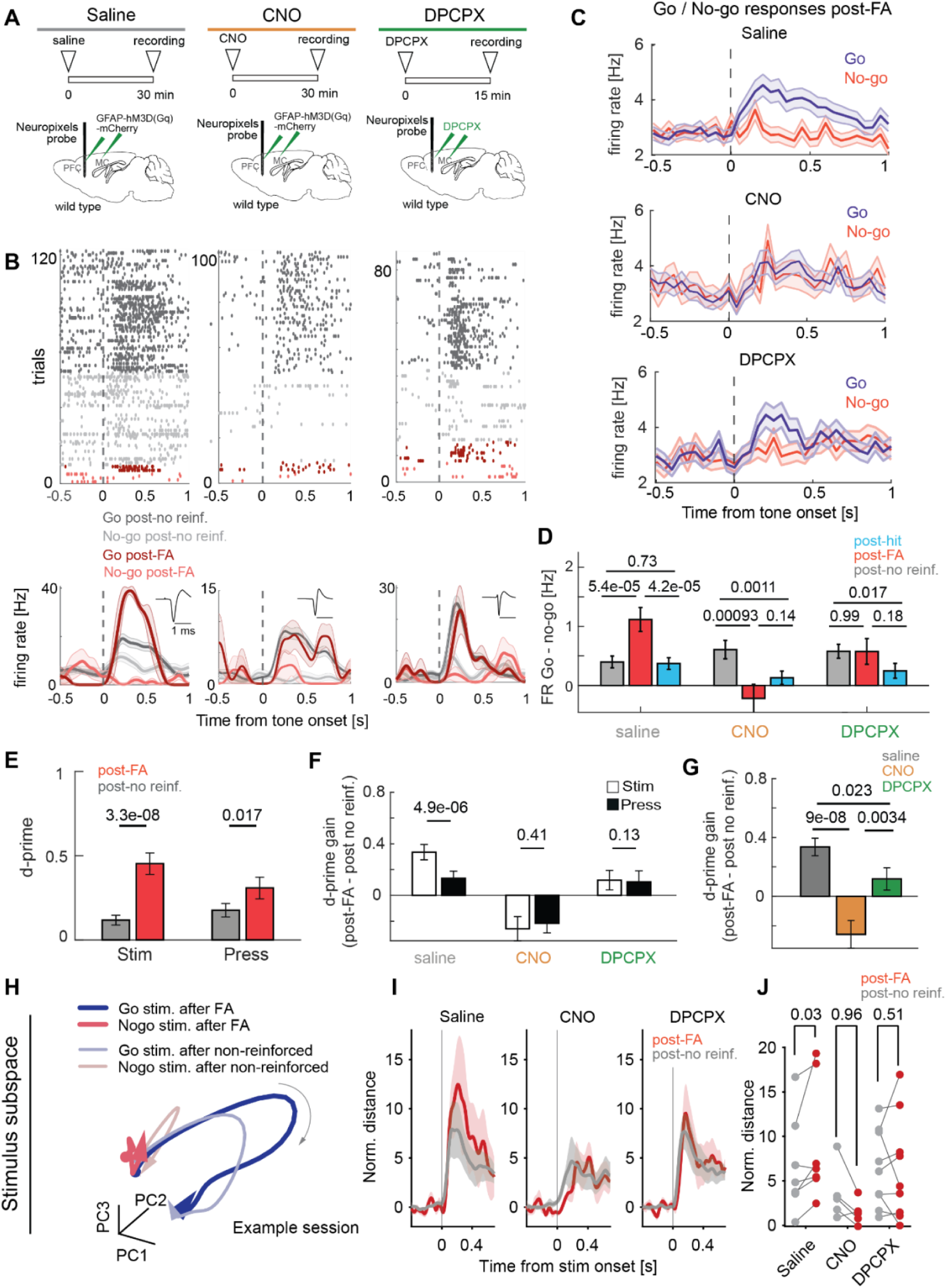
Astrocyte calcium manipulations and A1 receptor blockers occlude improvements in neuronal stimulus encoding post-false alarm. **(A)** Experimental design for saline (left), CNO (middle), and DPCPX (right) recordings. **(B)** Top: Raster plots showing activity on single trials for go and no-go stimuli after unreinforced (gray) and false alarm (red) trials for example units. Bottom: Average activity for go/no-go stimuli after unreinforced and false alarm trials. Insets show units waveforms. Experimental conditions corresponding to **A**. **(C)** Population averaged firing rate for go and no-go stimuli on trials after a false alarm in saline (top), CNO (center), and DPCPX (bottom) conditions. **(D)** Quantification of the difference between go and no-go firing rates on trials after hit (blue), false alarm (red), and unreinforced (gray) trials. **(E)** Stimulus (left) and press (right) neuronal d-prime after false alarm versus d-prime after unreinforced trials for saline recordings. **(F)** Comparison of stimulus and press neuronal d-prime gain between unreinforced and false alarm trials, for saline, CNO, and DPCPX sessions. **(G)** Quantification of stimulus d-prime gain between saline, CNO, and DPCPX sessions. n = 526 neurons (n = 6 mice); n = 360 neurons, CNO (n = 5 mice); n = 429 neurons, DPCPX (n = 5 mice). **(H)** Example neural population trajectories projected in the first three PCs of stimulus-related activity subspace for go and no-go stimuli after false alarm or unreinforced trials. The gray arrow indicates the progression of time. **(I)** Normalized Euclidean distance between go and no-go stimuli in saline, CNO and DPCPX session. **(J)** Comparison of normalized Euclidean distance between go and no-go stimuli trajectory after false alarm or unreinforced trials. n = 7 sessions, saline; n = 5 sessions, CNO; n = 9 sessions, DPCPX. P values show comparisons based on two-tailed paired t-test in **D-F**, on two-tailed unpaired t-test in **G**, and on one-tailed paired t-test in **J**. Data show mean ± SEM.

We thus hypothesized that astrocyte signaling could elicit the improved behavioral d-prime on trials after a false alarm by altering neuronal discriminability of go and no-go stimuli on these trials. Focusing on the epoch after stimulus tone onset, we analyzed the difference in single cell firing rates between go and no-go stimulus trials following specific trial outcomes. In saline sessions, we observed a clear firing rate difference between go and no-go stimuli after a false alarm, which was not apparent in CNO and DPCPX sessions (**Figures 4B and 4C**). This difference in go/no-go firing rates in trials following a false alarm was larger compared to other trial outcomes in saline sessions, but not in CNO and DPCPX sessions (**Figures 4B-4D**, **S14D, and S14E**). Thus, a post-false alarm air puff elicited a stimulus-specific change in firing rate on the following trial which did not occur when astrocyte calcium dynamics or A1 receptors were blocked. These findings indicate that astrocyte signaling specifically alters task-relevant neuronal encoding on trials following a surprising outcome rather than non-specifically suppressing neuronal spiking. To quantify the discriminability of go vs no-go stimuli in neuronal responses, we computed the neuronal stimulus d-prime between responses to these stimuli to show firing rate differences in units of trial-to-trial variability. In saline sessions, neuronal stimulus d-prime increased significantly following a false alarm compared to trials with no previous reinforcement (**Figure 4E**). A direct comparison of stimulus and press neuronal d-prime gain confirmed that false alarms specifically enhanced sensory stimulus encoding in prefrontal cortex, providing a potential substrate for increased behavioral stimulus sensitivity (**Figure 4F**). Such a specific gain in stimulus discriminability after a false alarm was absent in CNO and DPCPX sessions. The overall change in neuronal d-prime gain after a false alarm was also significantly larger in saline sessions compared to CNO and DPCPX sessions (**Figure 4G**).

Astrocyte processes are capable of contacting tens of thousands of synapses and astrocytes show sustained temporal dynamics during learned behavior; we thus hypothesized that astrocyte activity following a surprising outcome could alter large-scale neuronal population dynamics. To evaluate the effects of astrocyte dynamics on neuronal population encoding, we reduced the dimensionality of neuronal population activity during the stimulus presentation and identified a stimulus-related subspace that captured separable population activity between go and no-go trials. We then calculated the distance in this subspace between the trajectories for go and no-go stimuli after false alarm and unreinforced trials (**Figure 4H**). In the saline condition, the normalized distance between go and no-go stimuli was significantly increased after a false alarm trial, while CNO and DPCPX treatment blocked this effect (**Figures 4I and 4J**). We did not observe an increase in normalized distance between press and no-press trials in a press-related subspace, confirming that previous false alarms specifically increased stimulus discriminability in large neuronal populations (**Figures S14F-S14H**).

We next evaluated the effects of a surprising correct rejection reward on neuronal task encoding. Overall neuronal firing rates were not significantly affected by CNO or DPCPX (**Figure S16A)**. Neuronal firing rates specifically after a surprising correct rejection reward were also not altered in the saline and CNO condition but were affected by DPCPX (**Figure S16B**). Interestingly, we saw no difference in the discriminability between go and no-go stimuli following rewarded correct rejection trials, but we did observe a decrease in the discriminability between press and no-press compared with unrewarded correct rejections (**Figures S16C and S16D**). This decrease in press signals was abolished by either activation of astrocyte DREADDs or DPCPX treatment (**Figures S16C and S16D**). We next evaluated the neuronal population activity following a correct rejection reward in stimulus- and press-related subspaces. In line with our single cell analysis, we found no change in the separability of population representations of the go and no-go stimulus tones following correct rejection rewarded trials (**Figures S16E-S16G**). However, when we evaluated the population activity in the press subspace, we observed a significant decrease in the distance between press and no-press trials in the population activity (**Figures S16H-S16J**). Finally, CNO and DPCPX treatment blocked the decrease in separability for press and no-press activity (**Figures S16I and S16J**). Taken together, these data suggest that correct rejection reward trials reduce pressing behavior, or response bias, on the subsequent trial via astrocyte ATP/adenosine signaling to reduce neuronal press signals in the cortex. This behavior stands in contrast to behavior after a false alarm trial, which leads to increased hits and reduced false alarms, consistent with increased stimulus discriminability in neuronal responses.

Our findings thus demonstrate that following a false alarm, phasic NE release and subsequent astrocyte calcium and purinergic signaling alters neuronal responses to improve discriminability of go and no-go stimuli on the next trial at the single cell and population levels (**Figure 5**). This signaling is context-dependent, as astrocyte calcium and purinergic dynamics following a rewarded correct rejection, which reduces press probability on the next trial, reduces the separability of press vs no-press signaling in single neuron and neuronal population responses. The differential effects of astrocyte calcium signals on neuronal activity for different task outcomes are consistent with context-dependent reinforcement integration by astrocytes, which in turn differentially alters neuronal task encoding. These data indicate that NE-astrocyte signaling plays a significant and nuanced role in learned behaviors.

**Figure 5.**
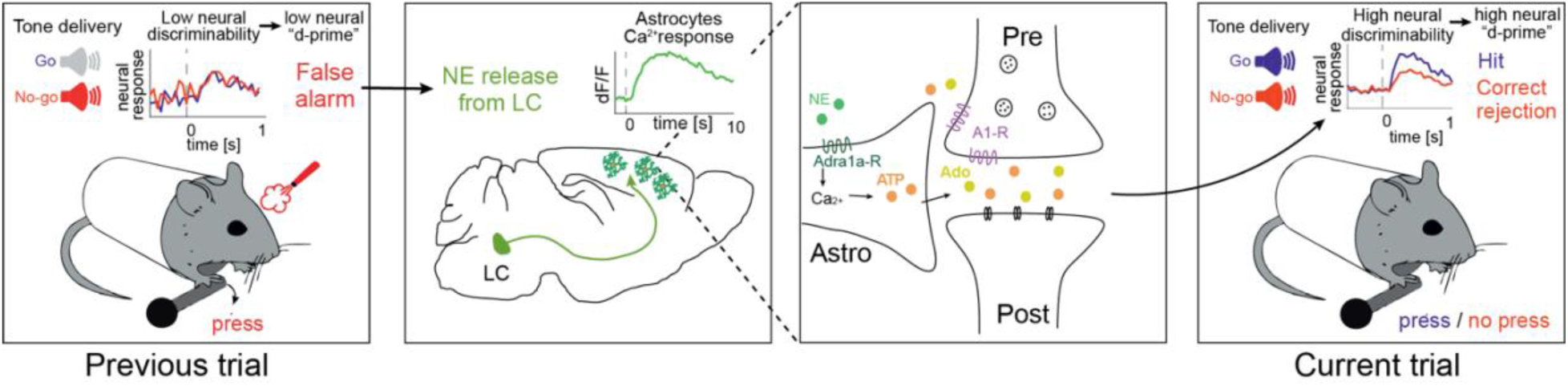
Model of the role of NE-astrocyte-adenosine signaling in learned behavior. Following a false alarm trial, LC-NE release leads to large and sustained calcium responses in cortical astrocytes via Adra1a receptors, which triggers A1 receptor activation on neurons to improve discriminability between go and no-go stimuli on the subsequent trial.

## Discussion

Using a behavioral task in which LC-NE activation facilitates serial, online learning, we demonstrate an instrumental role for astrocyte signaling in the acquisition and implementation of such learning in the brain. Transient NE release following an unexpected outcome acts directly on astrocytic Adra1a receptors to generate sustained calcium elevations, enabling a phasic NE reinforcement signal to bridge across trials, several seconds apart. These calcium dynamics are required for next-trial performance improvements and mediate neuronal response changes through extracellular ATP release and subsequent adenosine signaling via neuronal A1 receptors. Disrupting astrocyte calcium or blocking A1 receptors abolishes both the neuronal encoding changes and behavioral adaptation observed after a surprising outcome. Selective knockdown of Adra1a receptors on astrocytes blocks NE-driven calcium signal increases and reinforcement-driven behavioral improvement during task acquisition. Taken together, our findings demonstrate that astrocytes integrate NE-mediated prediction errors and propagate these signals across trials to alter neuronal task encoding and influence behavioral performance.

While NE has long been associated with arousal^8,10,11,21,55^ and global brain state regulation^20,84^, our behavioral experiments probe the role of phasic NE release on the timescale of hundreds of milliseconds, scaled by outcome expectation and thus characteristic of reinforcement learning signals. LC-NE activity and astrocyte calcium responses show a negative correlation with reward expectation, consistent with prediction error signaling; indeed, large NE and astrocyte calcium responses following a false alarm are consistent with signaling large reward prediction error. Importantly, astrocyte calcium signaling, but not changes in LFP power or neuronal population signals, persists across the inter-trial interval and predicts next trial performance. In addition, there is no relationship between changes in pupil diameter during the task and trial type or behavioral performance^7^, and others have similarly shown that phasic NE release associated with behavioral tasks engages mechanisms largely separable from those underlying arousal^15,32–34,36,85^. Finally, unexpected outcomes that both lead to large LC-NE and astrocyte calcium responses can have opposite effects on behavior – an increase in hits after a false alarm but a decrease in hits after a correct rejection reward. Thus, it is unlikely that our findings can be simply explained by a change in cortical state alone.

Astrocytes express high levels of α1-adrenergic receptors and are highly responsive to phasic NE release^45–50,86,87^. We propose that astrocytes act as temporal integrators of NE-mediated prediction errors: phasic NE is transformed into prolonged calcium signals that trigger ATP release and adenosine-dependent modulation of nearby synapses. While adenosine has been shown to act on A1 receptors on presynaptic terminals to inhibit excitatory transmission in slices^74,88,89^, it would be expected to have more complex effects *in vivo*^90^, by inhibiting excitatory transmission to both excitatory and inhibitory neurons, and thus affecting inhibitory transmission directly and indirectly^89,91^, as well as activating adenosine receptors on astrocytes themselves^92^. Our findings extend these previous studies to suggest that *in vivo,* during a complex behavior, astrocytic ATP/adenosine signaling alters task-specific neuronal encoding of key behavioral components. Our model is supported by recent analyses of NE signaling to astrocytes and subsequent purinergic synaptic modulation of neurons in the hippocampus *ex vivo*^74^, and by astrocytic purinergic regulation of swimming in zebrafish^75^.

Notably, astrocytes shape neuronal task encoding depending on the context and conditions of reinforcement: NE-induced calcium increases following a false alarm enhance stimulus discriminability in neuronal responses on the next trial, improving hit rates and reducing false alarms. In contrast, NE release following a correct rejection-reward results in astrocyte calcium elevations that lead to reduced press encoding, supporting an adaptive decrease in press probability. The neuronal changes occur without a generalized effect on firing rate, indicating selective modulation of population coding rather than nonspecific state changes^41,42^. Astrocyte heterogeneity may contribute to this contextual tuning, as we observe partially non-overlapping populations of astrocytes responsive to these task outcomes (**Figures S3F and S3G**). Indeed, the influence of distinct sets of synapses may be reflected in distinct profiles of astrocyte calcium responses under different activation conditions.

More broadly, our findings support emerging theoretical frameworks proposing that astrocytes contribute to neural computation by integrating signals over longer timescales and providing contextual guidance to neuronal networks^42,93,94^. By sustaining reinforcement signals long enough to influence subsequent behavior, astrocytes may be crucial more generally for constructing and refining internal models of the world—a fundamental component of cognition—at time scales consequential for natural behavior.

## Experimental Model

### Mice

All procedures performed in this study were in accordance with the Massachusetts Institute of Technology’s Animal Care and Use Committee and the Guide for the Care and Use of Laboratory Animals published by the National Institutes of Health. Male and female mice more than 2 months old were used in this study. Mice were kept in a room with a (12:12) reversed light/dark cycle with controlled temperature and ventilation (20–22 °C; 40–60% humidity). All experiments were performed during the dark period of the cycle. The *Dbh-cre* line (B6.FVB(Cg)-Tg(Dbh-cre)KH212Gsat/Mmucd, MMRRC) was used for specific expression of Gi-DREADD virus in norepinephrine-expressing neurons of the locus coeruleus (LC). The *Adra1a^fl/fl^* line (University of California San Francisco) was used for astrocyte-specific Adra1a knock-down experiments. C57BL/6 wild-type mice (Jackson Laboratories) were used for all other experiments.

## Method Details

### Viral vectors

For dual imaging of cytosolic astrocyte calcium activity and neuronal calcium activity, wild-type mice were injected with AAV5-gfaABC1D-cyto-GCaMP6 (Addgene #52925-AAV5) and AAV1-Syn-NES-jRGECO1a-WPRE (Addgene #100854-AAV1) into motor cortex (MC). For chemogenetic manipulation of astrocytes, wild-type mice were injected with AAV5-GFAP-hM3D(Gq)-mCherry (Addgene #50478-AAV5) into motor cortex (MC) and prefrontal cortex (PFC). For chemogenetic silencing of the LC while imaging in the MC, *Dbh-cre* mice were injected with AAV5-hSyn-DIO-hM4D(Gi)-mCherry (Addgene #44362-AAV5) into the LC and AAV5-gfaABC1D-cyto-GCaMP6 (Addgene #52925-AAV5) into the MC. For Adra1a knock-down experiments while imaging in the MC, *Adra1a^fl/fl^* mice were injected with AAV5-GFAP(0.7)-mCherry-T2A-iCre (Vector Biolabs #VB1132) and AAV5-gfaABC1D-cyto-GCaMP6 (Addgene #52925-AAV5) into the MC. For LC axonal stimulation, *Dbh-cre* mice were injected with AAV1-EF1a-double floxed-hChR2(H134R)-EYFP (Addgene #20298-AAV1) in the LC and AAV5-gfaABC1D-lck-GCaMP6 (Addgene #52924-AAV5) into the visual cortex. For GRAB sensor imaging, wild-type mice were injected with AAV9-hSyn-mRuby3-NE2h (WZ Biosciences) or with AAV9-hSyn-GRAB-ATP1.0 (Addgene #167577; viral packaging done in-house) into the MC.

### Stereotactic surgery

Surgeries were performed aseptically under isoflurane anesthesia while maintaining body temperature at 37.5°C on a heating pad (ATC2000, World Precision Instruments). Mice were given pre-operative slow-release buprenorphine (1mg/kg, subcutaneous injection) as well as pre- and post-operative meloxicam (5mg/kg, subcutaneous injection). Mice were placed in a stereotaxic frame (model 51725D, Stoelting), scalp hair was removed with a depilatory cream, and the incision site was sterilized using betadine and 70% ethanol. The skull was exposed and the conjunctive tissue was removed using hydrogen peroxide. The skull was positioned such that the bregma and lambda marks were aligned on the anteroposterior and dorsoventral axes. For virus delivery, a small craniotomy (0.5 mm) was drilled above the region of interest and virus was injected using a glass pipette with a 50 µm diameter tip. For delivering Gi-DREADD virus to the LC, 300 nl of virus was injected (rate: 50nl/min). Coordinates for targeting the LC were (in mm): -5.2 to -5.0 anterior-posterior (AP), ±0.9 medial-lateral (ML), -2.8 dorsal-ventral (DV). For delivering Gq-DREADD virus to the cortex, 200 nl of virus was injected (rate: 50nl/min) into the motor cortex (MC) and prefrontal cortex (PFC) bilaterally. Coordinates for targeting the PFC were (in mm): +2.0 AP, ± 0.3 ML, -0.5 DV. Coordinates for targeting the MC were (in mm): +0.5 AP, ±1.5 ML, -0.3 DV. For two-photon imaging, a round 3 mm diameter craniotomy was drilled over the left MC. Three to four injections of 250nl of gfaABC1D-GCaMP6 and syn-JRGECO1a virus were made into layer 2/3 of the MC (DV -0.3mm, rate: 50nl/min), waiting 5 minutes before withdrawing the glass pipette after each injection to avoid backflow. For delivering GRAB-NE sensors, a total of 400-500nl of hSyn-mRuby3-NE2h virus was injected in four locations of the MC craniotomy. For delivering GRAB-ATP sensors, a total of 500-600nl of hSyn-GRAB-ATP1.0 was injected in three locations of the MC craniotomy. A cranial window, made of two 3 mm coverslips centered on a 5 mm coverslip (CS-3R, CS-5R; Warner Instruments) and glued together with ultraviolet adhesive, was positioned over the craniotomy and attached to the skull using dental cement (C&B Metabond, Parkell). To avoid light reflection and absorption, the dental cement was mixed with black ink pigment (Black Iron Oxide 18727, Schmincke). A head plate was then affixed to the skull with dental cement for head fixation during the behavioral task. Following surgery, animals were allowed to recover on a heating pad. Their recovery was monitored for a minimum of 72 hours and anti-inflammatory (Meloxicam) injections were provided as needed. Animals recovered for at least 5 days before starting water restriction for behavioral experiments.

### Behavior set-up

Mice were head-fixed on a behavior rig and confined in a polypropylene tube to limit body movements. Their left forepaw was able to move a lever built with a 1/16-mm-thick brass rod attached to a piezoelectric flexible force transducer (LCL-005, Omega Engineering), and a metallic lick spout placed near the mouse’s mouth and connected to a custom-made 80 lick detector was used to deliver water rewards (∼5 μL drop of water). A small tube, pointing towards the mouse’s facial area and at a distance of 3 cm, was used to deliver air puff punishment (compressed air at 40 psi for 0.3s). Voltage signals from the transducer were recorded through a microcontroller board (Arduino UNO Rev3). Voltage signal from the transducer was converted to lever movement in degrees using calibration data from video analysis. A second microcontroller board was used to control a 5 mm yellow LED light placed 8 cm in front of the mouse and two solenoid valves (Parker #003-0141-900) for water and air puff delivery. 4 or 12 kHz sound stimuli of 0.5s duration were delivered using a single speaker located at a distance of 30 cm from the mouse. The speaker frequency range was calibrated using a USB calibrated measurement microphone (UMIK-1, Mini DSP) and the Room EQ Wizard software, and the sound stimulus intensities were established by a sound level meter. Two behavior rigs were used: one for general behavior and electrophysiology, and one for two-photon imaging. Noise levels were comparable across both rigs (in dB with Z noise frequency weightings): 7.8±1.1, 8.8±1.0, 14.3±0.8, and 14.7±0.9 for 4 kHz; and -4.0±1.2, - 1.7±1.1, -1.9±0.9, and 0.3±0.7 for 12 kHz. The behavior rig was connected to a computer running a custom written MATLAB (Mathworks) script that was able to record lever voltage, while controlling the timing of light cue, sound (using Psychtoolbox), water, and reward. Behavior rigs were assembled primarily with optomechanical components (Thorlabs).

### Behavior task

Upon recovery from surgical procedure, mice were gradually put on a water restriction schedule, eventually receiving 1-1.5 mL of water per day. Body weight was maintained above 90% of their pre-restriction weight. Mice were trained to hold still for 1.5s during the cue period (LED on), to wait for a delay to hear a tone, and to either push the lever to obtain a reward or to refrain from pushing to avoid a punishment. Mice learned to push the lever when they heard a go tone (12 kHz frequency) and hold still when they heard a no-go tone (4 kHz). After the onset of the 0.5s sound stimulus, mice had 0.8s to respond or hold still. If they pressed the lever during a go trial, they received a water reward. If they pressed during a no-go trial, they received a mild air puff punishment. Absence of response on go trials (miss) or not pressing the lever during no-go trials (correct rejections) were not reinforced. To vary the level of stimulus evidence, 4 intensities were used per frequency for a total of 8 different stimuli. Tone intensities used were 5, 15, 25, and 35 dB. These values were calculated by measuring the sound pressure level for either go or no-go frequency and subtracting the noise level of that given frequency. A lever press (hit or false alarm) was determined when the lever position reached a threshold value of 3 to 4° (depending on the animal) from the position at the beginning of the trial, while absence of a lever press (miss or correct rejection) was determined if the lever absolute position stayed below a value of 2.2°. Premature lever presses, which occurred in the delay period between light cue off and tone onset, were considered early presses and aborted the trial. The delay between light cue off and tone onset was randomized following a gaussian distribution (mean: 0.65s; standard deviation: 0.15s). Trial order was pseudo-randomized to ensure that the same amount of go or no-go trials were presented every fourth trial, and that each tone intensity was presented every eighth trial. Each trial was followed by a 2.5s reinforcement consumption period and 1.5s cue period (LED on). Mice were taken through two stages of training until they became proficient at the task. During the first phase of training, mice learned to associate a lever press with reward and to detect a go tone. In this phase, only go tones (tone: 12 kHz at 35 dB for 0.5s) were used. The same trial sequence as in the full task was used, but with an extended duration of the response window (30s instead of 0.8s). The animal was switched to the next stage once they made more than 80% of lever presses for 50 consecutive trials within a period of 0.8s after tone onset. During the second phase of training, no-go trials (tone: 4 kHz at 35 dB for 0.5s) were introduced and the response window was reduced to 0.8s after tone onset. Mice used for the Adra1a KD experiments were evaluated in this second stage of training, upon the introduction of no-go tones. Training in the second phase lasted until mouse performance reached 85% hit rate and less than 30% false alarm rate for two consecutive sessions. The last stage was considered the full task, in which 4 intensities of each tone were introduced (5, 15, 25, and 35 dB). For physiological recordings, a 0.25s delay between the timing of lever press and reward/punishment was introduced for neuronal and astrocytic calcium imaging and 0.8s delay for GRAB sensor imaging. For correct rejection with surprising reward experiments, mice received a water reward randomly on a quarter of correct rejection trials on a subset of recording sessions. For ATP imaging, mice were imaged at the second stage of training (go and no-go trials with 1 intensity each).

### Two-photon calcium imaging

After training to proficiency, awake mice that were previously implanted with a cranial window were head-fixed, and the fluorescent sensors (GCaMP, jRGECO, mCherry, GRAB-NE, and GRAB-ATP) were imaged using galvo-galvo scanning with a Prairie Ultima IV two-photon microscopy system (Bruker). Two-photon excitation was provided by an insight tunable laser (InSight X3+ or MaiTai, Spectra-Physics). Imaging of GCaMP was done at a wavelength of 920 nm, imaging of mCherry was done at 1020 nm, dual-imaging of GCaMP and jRGECO was done at 1000 nm, and imaging of GRAB-ATP was done at 920nm. A 16x 0.8 numerical aperture objective (Nikon) and 2-4x optical zoom was used for imaging, with a frame rate of 5-6 Hz for GCaMP imaging and a frame rate of 16 Hz for dual imaging of GCaMP and jRGECO or GRAB-ATP. A voltage signal indicating the start of each trial was recorded by the system for alignment with the behavior (Bruker). Imaging sessions lasted ∼25 minutes per animal. GRAB-NE imaging was done at 920nm with a Prairie Ultima IV two-photon microscope attached to an XLUMPlanFLN 20x 1.00 NA (Olympus) objective with a frame rate of 30 Hz.

### LC axonal stimulation

Awake mice that were previously implanted with a cranial window were headfixed and ChR2 expressing axons were stimulated via a 473 nm laser through the objective for 0.5 – 2 seconds at 30 Hz. During stimulation, the shutter was closed and no images were captured.

### Neuropixels recordings

After training to proficiency, mice that were previously injected with Gq-DREADD virus were anesthetized with isoflurane. The silicone elastomer and gel foam (Pfizer) covering the skull were removed and a 500 μm diameter craniotomy was performed over the MC (+0.5 mm AP; ±1.5 mm ML) and PFC (+2 mm AP; ±0.3 mm ML). The dura was punctured and the craniotomy was protected with saline and a piece of gel foam (Pfizer). The skull was covered again with silicone elastomer and the mouse was allowed to recover for one day before recording. The next day, the awake animal was head-fixed and the silicone elastomer and gel foam were gently removed. A 0.9% NaCl solution was used to keep the surface of the brain wet for the duration of recordings. After placing the animal on the recording set up, a reference silver wire was submerged in the saline solution on the skull surface. The neuropixels probe was then referenced on bregma and the surface of the brain. The probe was slowly lowered (1 min per mm) using a motorized micromanipulator (MP-285, Sutter Instrument Company) until the desired depth (∼2000 µm) was reached. The probe was left to sit in the brain for at least 15 minutes prior to initiating the recording to minimize probe drift during the recording. Time stamps of each trial start were recorded by the neuropixels Spike GLX software. Physiology data was collected at a rate of 30 kHz using the Spike GLX software. Spike sorting was done using kilosort, and spikes were manually curated using Phy to remove artifacts picked by the algorithm such as ill-shaped units, units with low amplitude spikes, units with low spike rate, or units without a clear refractory period. Spike times were verified with cross correlograms to combine units or eliminate duplicates. For each unit, parts of the recordings with obvious drift (unit firing rate abruptly decreasing) were excluded. At the end of each session, the probe was coated with DiI, DiO, or DiD and reinserted at the same location and depth to mark the recording track. The craniotomy was covered again with gel-foam and silicone elastomer to allow recording on the next day. After the last recording, the brain was collected (see methods for histology) and imaged on a confocal microscope (Leica SP8). Slices with DiI, DiO, or DiD probe tracks were aligned to the Allen Brain Atlas using SHARP-Track^95^ to verify the position of the probe.

### Saline/CNO behavior

Clozapine-N-oxide dihydrochloride (CNO; Hello Bio) was dissolved in saline injectable sterile solution (0.9% sodium chloride) and administered at a low concentration of 0.2 mg/kg^62,96^. The CNO solution or saline control was injected intraperitoneally 30 minutes before each behavior session.

### Saline/DPCPX behavior

8-cyclopentyl-1,3-dipropylxanthine (DPCPX; MilliporeSigma) was dissolved in DMSO and saline, and administered at a concentration of 10 µM^79^. Gel foam soaked in the 10 µM DPCPX solution or saline control was applied directly on the skull of a head-fixed animal that previously underwent craniotomies on the MC and PFC (see methods for neuropixels recording). The behavior session was started 15 minutes after initial DPCPX or saline application, allowing time for the solution to penetrate through the previously-punctured dura.

### Fluorescence in situ hybridization

Mice were deeply anesthetized with isoflurane and transcardially perfused with ice cold phosphate buffered saline. The brain was then rapidly dissected and flash frozen in a cryomold held above liquid nitrogen. The frozen tissue was embedded in Tissue-Tek OCT compound (Sakura Finetek) prior to slicing on a cryostat (Leica CM 3050S). 14 µm sagittal section were mounted directly on SuperFrost Plus slides (VWR) and stored at -80°C. Fluorescence in situ hybridization was conducted using the manufacturer’s recommended protocol and reagents (RNAscope Multiplex Fluorescent V2 Assay, Advanced Cell Diagnostics). The probes used were against *Adra1a* (408661, Advanced Cell Diagnostics), *mCherry* (431201-C2, Advanced Cell Diagnostics), and *Slc1a3* (430781-C4, Advanced Cell Diagnostics) and were labelled with TSA Vivid dyes 570 (323272, Advanced Cell Diagnostics), 520 (323271, Advanced Cell Diagnostics), and 650 (323273, Advanced Cell Diagnostics) respectively. Labelled sections were stained with DAPI, mounted with Prolong Gold Antifade mounting media (ThermoFisher Scientific), and imaged on a confocal microscope (Leica SP8) using a 20x objective. Analysis was performed using QuPath^97^ to estimate the number of puncta of each labelled RNA per cell. The difference in *Adra1a* expression between groups of cells was determined via a mixed effects ANOVA corrected for multiple comparisons. Mouse of origin for each cell was used as the random effect. Only one section was included per mouse in the analysis.

### Histology

Mice were deeply anesthetized and transcardially perfused with cold 0.1 M phosphate buffered saline followed by 4% paraformaldehyde (PFA). Brains were collected and post-fixed in 4% PFA overnight at 4°C. Brains were then sectioned into 100 μm coronal sections with a vibratome (Leica VT 1200S). Sections were mounted with Vectashield mounting media (Vector Laboratories) and imaged with a confocal microscope (Leica SP8) using a 10x objective to confirm viral expression in target regions.

### Glial cell culture and adenosine assay

Brains of two-day-old neonatal C57BL/6 mice were removed after decapitation and cortices were dissected out using a dissecting microscope. The tissue was dissociated to a single cell suspension using enzymatic and mechanical trituration. The collected cell suspension was cultured in 35 mm culture dishes in Astrocyte medium (ScienCell) containing 10% fetal bovine serum and was maintained at 37°C in a humidified atmosphere containing 5% CO_2_. The medium was changed every 3–4 days. Cultures were used for the adenosine assay after 14 days. For the adenosine assay, the cell medium was replaced with physiological saline containing 120 mM NaCl, 4 mM KCl, 1.2 mM MgSO_4_, 10 mM glucose, 2 mM CaCl_2_, 10 mM HEPES, pH 7.35. During the experiment, cells were incubated with 20 μM NE for 5 min and an aliquot of saline was used to measure adenosine using an adenosine assay kit (Abcam) and a microplate reader (Perkin Elmer Enspire) after 30 min.

For evaluating effects of alpha1 adrenergic receptor inhibition, cells were pre-incubated in saline containing 10 µM prazosin hydrochloride (Sigma) for at least two hours before NE treatment.

### Behavioral analyses

For each animal, the average press probability for each stimulus tone was computed and averaged across no-go and go tones to determine the false alarm rate (*F*) and hit rate (*H*), respectively. Behavioral stimulus discriminability was then quantified by calculating the d-prime (*d*′) as follows:

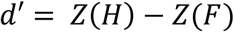

where *Z* represents the z-score. Behavioral bias, reflecting the tendency to press regardless of the stimulus identity, was quantified as follows:

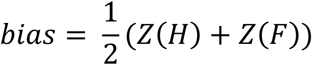

A value of 0 indicates no bias. A positive (negative) value indicates a press-oriented (withhold-oriented) bias. We calculated *H*, *F*, *d*′ and *bias* after specific trial outcomes and on surrogate data generated by shuffling the trial sequence (N=1,000 shuffles). We then subtracted the average shuffled *H*, *F*, *d*^′^, and *bias* from the values computed after a specific outcome to evaluate the effect of specific previous outcomes on these metrics, compared to their values computed irrespective of the previous trial outcome. We excluded parts of a session where the hit rate was lower than 40% and false alarm rate was higher than 50%, calculated using a 50-trial averaging window. For experiments with rewarded correct rejections, which have lower number of sessions per animal, we computed these metrics using all available trials. For these analyses we only included animals with at least two behavioral session and discarded 2 sessions with below chance level performance. Trials with presses occurring before the stimulus presentation (early presses) were removed from all analyses.

### Imaging data analysis

Time-lapse imaging sequences were corrected for x and y movement using the template-matching ImageJ plugin. For neuronal analysis, regions of interests (ROI) were automatically identified using Suite2P, manually curated, and the fluorescence intensity in time for each ROI was averaged. For astrocyte analysis, ROIs were manually drawn using ImageJ’s ROI manager tool, and fluorescence intensity over time was measured with ImageJ’s multi-measure feature. Baseline fluorescence *F*_0_ was computed as the average raw fluorescence of each cell over the entire linearly detrended recording session. Relative changes in fluorescence Δ*F*/*F*_0_ (%) were calculated as the raw fluorescence at each time point minus *F*_0_, divided by *F*_0_, and multiplied by 100. Single-trial, tone-aligned ΔF/F₀ responses were downsampled to the minimum sampling frequency within each experimental condition using spline interpolation, enabling pooling of responses across ROIs. For rewarded correct rejection experiments, data from one animal exhibiting strong negative deflections in calcium traces were curated by excluding trials with any ROI showing a negative deflection exceeding 2 standard deviations across the entire trace. Two sessions with fewer than 25 trials post-curation were excluded from further analysis. We quantified the mean activity and response timescale of each cell for specific trial types (hit, miss, correct rejection, and false alarm).

Peak activity was calculated as the maximum Δ*F*/*F*_0_ within 0-6.27 s seconds after tone onset, where 6.27 s corresponds to the median time between consecutive tones. To quantify response duration, we first identified time points at which the mean activity of each cell across specific trial types deviated positively or negatively from 0 by more than one standard deviation of the baseline activity variability (from -1 to -0.2 seconds from tone onset), computed across all trials. We then clustered consecutive threshold-exceeding time points (separately for positive and negative deviations) within 10 seconds following tone onset. Cells were classified as responsive if they exhibited at least one activity cluster, and as responsive until the next trial if the cluster offset exceeded 6.27 s (median next tone onset). The response timescale of each responsive cell was quantified as the duration of the longest activity cluster. Correlation of post-FA astrocyte activity and next-trial behavioral success was computed as the across-session Pearson correlation between (i) the session-wise mean astrocyte response in FA trials (cell-averaged, then time-averaged [-1,0]s from the minimum next tone onset and z-scored across all trials) and (ii) the sessions’ mean success rate on trials following a FA. A similar pipeline was used to compute correlation values between all signals in Table 1 and next-trial success probability. To identify temporal patterns in astrocyte responses across different trial types, we performed hierarchical clustering on z-scored, baseline-subtracted, and concatenated average Δ*F*/*F*_0_ traces from tone onset to median next tone onset. For clustering analyses, response duration thresholds were computed as one standard deviation of the concatenated responses. Agglomerative hierarchical clustering was applied to Euclidean dissimilarity matrices using Ward linkage for cluster grouping. The optimal number of clusters was determined by maximizing the Calinski-Harabasz index. For comparability between response modulation on hit and FA trials, z-scored, baseline-corrected signals were used to compute the slope of calcium responses as a function of tone intensity. Astrocytes were divided into low and high FA-responsive astrocytes based on a median split of peak FA responses. For analysis of GRAB sensors, following motion correction, responsive ROIs were manually annotated in ImageJ and baseline correction was performed using the mean fluorescence of the entire session to generate a Δ*F*/*F*_0_ signal. The peak amplitude and Area Under the Curve (AUC) values of the peaks following reinforcement was calculated within a specified time window, [0 3]s for GRAB_NE2h_ and [0 1.5] s for GRAB_ATP_, from the signals aligned to the time of reinforcement. Statistical tests were performed using GraphPad Prism version 10.0.0 for Windows for GRAB_NE2h_ data.

### Neuropixels data analysis

For each neuron, we calculated the mean (*μ*) and standard deviation (*σ*) of the firing rate in sliding windows 50 ms long computed over go or no-go stimulus presentations and over press or no-press response, selecting trials after specific trial outcomes. All time-resolved single-cell metrics were averaged in the 0.05–0.25 s post-tone onset for quantitative comparisons. The stimulus and press neural *d*′ (a measure of target task variable *T* discriminability based on the activity of an individual neuron) in each time window was then computed as follows:

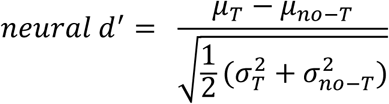

where *T* = *go* for stimulus d-prime and *T* = *press* for press d-prime, respectively. We set neural *d*^′^ values to zero in rare cases when a neuron did not fire any spike across all trials to both go and no-go stimuli or press and no-press response. We also excluded the rare cases when the neuron fired some spikes but with no variation of firing rate across-trials within both go and no-go conditions. To assess firing rate changes after specific outcomes, we compared observed rates against surrogate data (median of N = 5000 shuffles, stratified by current trial type). Significant *d*^′^differences reported in the paper were all successfully validated against shuffled distributions (not shown). One session was excluded from all Neuropixel analyses due to a low number of trials (n = 23) after removing early press trials.

For state-space analysis, we used neurons that were recorded in the sessions that included at least one trial for target trial type. We took population vectors (the z-scored activity of each neuron at each 10 ms time bins) in the window [-0.4 0.7]s around stimulus onset, we concatenated population vectors for the go and no-go stimulus trials or press and no-press response trials to define stimulus and press subspace, respectively. Principal Component Analysis (PCA) was used to the population vectors within the [0 0.25] s from stimulus onset to reduce the dimensionality of the population, retaining the number of principal components (PCs) that explained 80% of the total variance of the population vector dataset. To statistically evaluate the difference of each neuronal trajectory at each time point across trial types in the PC space, we projected the data from a random subset of trials (50%) onto each axis. The projection of each axis was computed as the dot product. The procedure was repeated 500 times. We then computed the Euclidean distance between trajectories of target trial types. To normalize these distances, we computed *D*_*norm*_ = (*D* − *μ*_*baseline*_)/*σ*_*baseline*_, where *μ*_*baseline*_ and *σ*_*baseline*_ are the mean and standard deviation, respectively, of the distances calculated during the pre-stimulus window [-0.4, 0] s. For comparison, distances were averaged over the [0.05 0.25] s from stimulus onset. For population decoding analyses, we trained a linear support vector machine to distinguish between FA and non-FA trials using the same z-scored population vectors employed in the state-space analyses. To balance the training set, non-FA trials were randomly resampled 50 times to match the number of FA trials. In each resampling, 25 neurons were randomly selected to control for population size differences. The decoder was trained with 2-fold cross-validation across trials and used lasso regularization, with the regularization parameter selected via 2-fold cross-validation.

We used the fieldtrip toolbox^98^ for Local Field Potential (LFP) power analysis. Raw digitized neurophysiological signals were converted to voltage. Channels exhibiting aberrant noise were excluded by thresholding the median absolute deviation (MAD) of each channel’s temporal standard deviation (5 MAD from the median). Data were bandpass-filtered (1–300 Hz, 4th-order Butterworth, two-pass) and notch-filtered at 60 Hz harmonics to remove line noise. Signals were downsampled to 1,000 Hz and re-referenced to the common median. PCA was used to reduce the dimensionality across channels, retaining components explaining at least 90% of variance for subsequent analyses. Time-frequency representations (TFR) were computed for each PC from reinforcement-aligned single-trial data using the multitaper method with a 0.5 s sliding window and 4 Hz half-bandwidth spectral smoothing (3 DPSS tapers in total). Median reaction time plus 0.25s was used to align correct rejection and miss trials. Power changes relative to baseline were computed for individual trials as the log-transform of TFR values minus their average baseline values ([-3 -0.5] s). Power values were first averaged across trials corresponding to a specific outcome, and then across PCs, to compute power changes for each session. Gamma power was computed as the average power in the [30,80] Hz band. Peak power change was computed as the maximum between 0 and minimum time of next cue, power values before next cue onset as the mean 3.5-4 s after reinforcement.

### Pupillometry

A high-speed ThorLabs camera recorded facial and pupil dynamics during head-fixed behavior, and the resulting videos were processed post hoc with DeepLabCut to extract eight xy-coordinates surrounding the pupil. ∼500 randomly selected frames were manually labeled (∼20 frames/ video) and a resnet v1 50-based convolutional neural network was trained to predict the location of the 8 markers (4 diametrically opposite pairs) around the circumference of the pupil for 50,000 iterations. Once the network was trained, it was used to automatically label pupil coordinates on all videos, and the quality of the labeling was manually evaluated by observing at least 10 labeled videos. The network was refined with additional frames for each experimental cohort. Linear interpolation was used to fill in missing values due to blinking. Pupil diameter was then computed for each of the 4 xy-coordinate pairs, averaged across all 4 diameters and z-scored to yield a stable, session-wide measure. Any major outliers in label position (a label suddenly jumping away from the eye) were removed post hoc by removing values above the 99th percentile of pupil values. Correlations between pupil diameter changes in FA trials and next-trial behavioral success were computed from reinforcement-aligned pupil traces following the same procedure described in the *Imaging data analysis* section.

### Statistics

Throughout the paper we used Student’s t-statistics to compute P values of one-sample, paired, and unpaired comparisons, apart from the cases listed below. For neural population trajectory distance comparisons, one-tailed t-statistics was used to confirm single-cell level effects. For correlation with next-trial behavior, one-tailed t-statistics was used to identify whether signals in the direction evoked by FA were predictive of next trial behavioral success. For Adra1a receptor KD experiments, we used a one-way ANOVA to compare astrocyte responses across trial types. For firing rate comparisons, which exhibit highly skewed distributions^99^, we employed nonparametric two-tailed Wilcoxon signed-rank or Kruskal-Wallis tests. Binomial test under the normal approximation was used to compare fractions of responsive cells. Multiple comparisons within the same dataset were corrected for family-wise error rate using the Bonferroni-Holm procedure. Significant time-frequency LFP power level deviations from baseline (two-tailed t-test) or across conditions (one-way ANOVA) were corrected with cluster analysis^100^. Surrogate data for cluster-based permutation test were generated via sign-flip within conditions (baseline comparisons) or shuffling condition labels (across-condition comparisons) at the group level, p < 0.05 was used for both the cluster-forming threshold and cluster-level statistical significance.

## Acknowledgments

We thank Taylor Johns for lab management and members of the Sur laboratory for many discussions and comments. We are grateful to James Schummers for technical advice, Juan Padilla for imaging, histology, and data pre-processing and Alexandria Barlowe for histology and animal colony management. This work was supported by NIH grants R01MH126351; R01MH133066; R01NS130361; R01MH085802; MURI Grant W911NF2110328; The Picower Institute Innovation Fund; and the Simons Foundation Autism Research Initiative through the Simons Center for the Social Brain (MS); NIH Fellowship F31MH129112 (GTD); European Union’s Horizon research and innovation programme grant agreement No. 101206609, Marie Sklodowska-Curie Global Fellowship - AstroError (MC); Japan Society for the Promotion of Science and Uehara Memorial Research Fund (YO); the JPB Foundation Postdoctoral Award (GF); and NIH Brain Initiative grant R01NS108410 (SP).

## Author contributions

Conceptualization: GTD, JS, MS; Methodology: GTD, AN, MC, JS, JP, PO, YO, PCS, SP, MS; Investigation: GTD, AN, JS, PO, JP, GF, GOS, KRJ, VB-P, TO, PCS; Visualization: GTD, MC, JP, GF, PO; Funding acquisition: MS, SP; Supervision: MS, SP; Writing – original draft: GTD, MS; Writing – review & editing: GTD, AN, MC, JP, KRJ, MS, SP.

## Data and materials availability

All data, code, and materials used in the analysis will be made available upon acceptance of the manuscript.

## Supplemental Figures and Table

**Figure S1.**
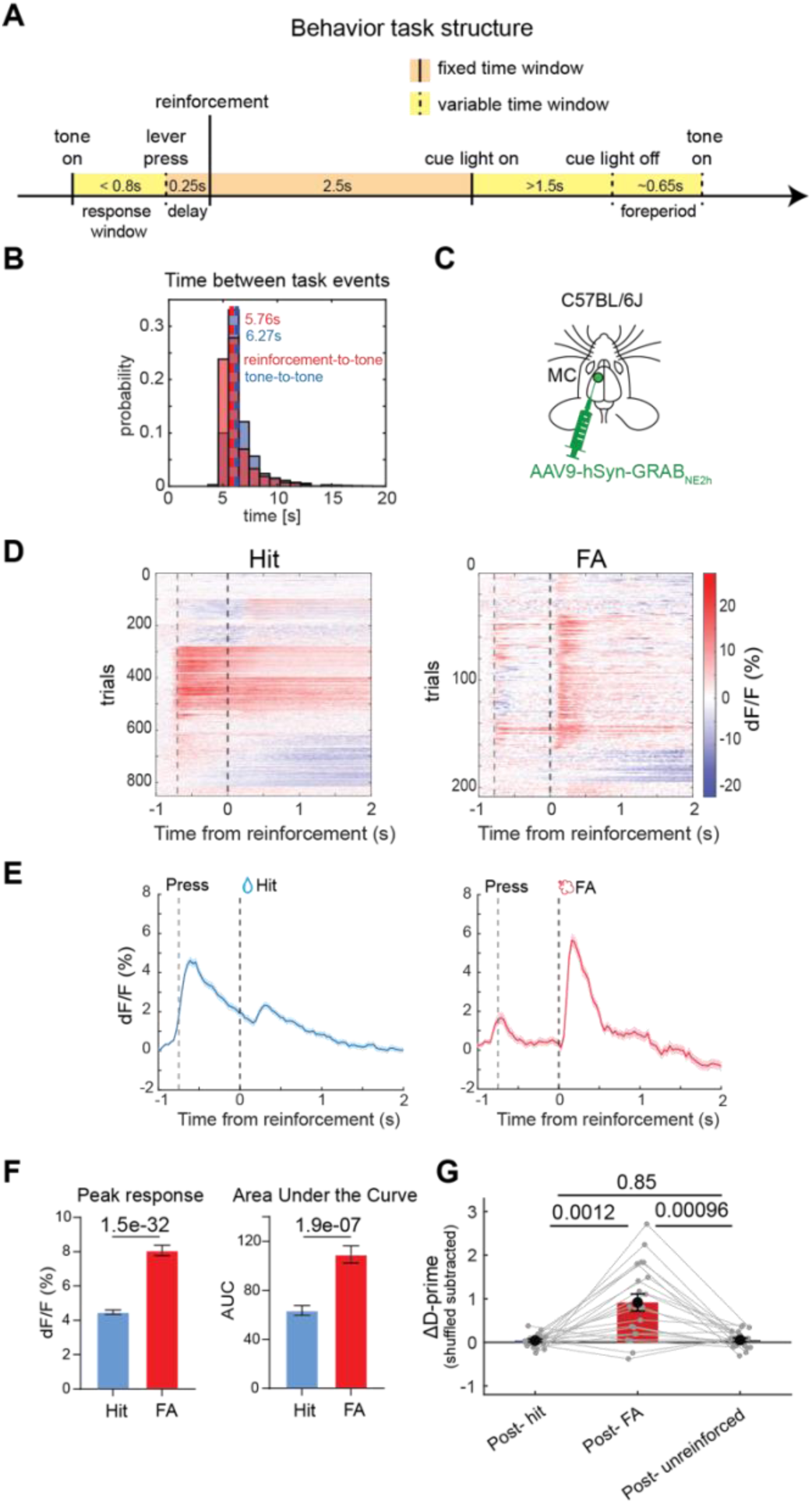
Task structure, in vivo characterization of NE dynamics, and d-prime changes after different trial types. **(A)** Depiction of behavioral task single-trial structure. **(B)** Across-trials distributions of time between tone-to-tone onset and reinforcement-to-tone onset with median values displayed (trials pooled across n = 21 mice). **(C)** Schematic for GRAB_NE2h_ labelling in the motor cortex. **(D)** Mean GRAB_NE2h_ dF/F signal on hit and false alarm trials, aligned to the time of reinforcement. **(E)** Mean GRAB_NE2h_ dF/F responses across individual hit and false alarm trials aligned to the time of reinforcement. Solid lines and shaded areas display mean ± SEM. Dashed lines denote the time of lever push and reinforcement. **(F)** Comparison of post-reinforcement peak amplitude (left) and AUC (right) of GRAB_NE2h_ responses. Unpaired t-test. Hits: n = 851 trials from 3 mice. FA: n = 208 trials from 3 mice. **(G)** Change in d-prime after hit, false alarm, and unreinforced trials (n = 20 mice). P values show comparisons based on two-tailed paired t-test. Data show mean ± SEM.

**Figure S2.**
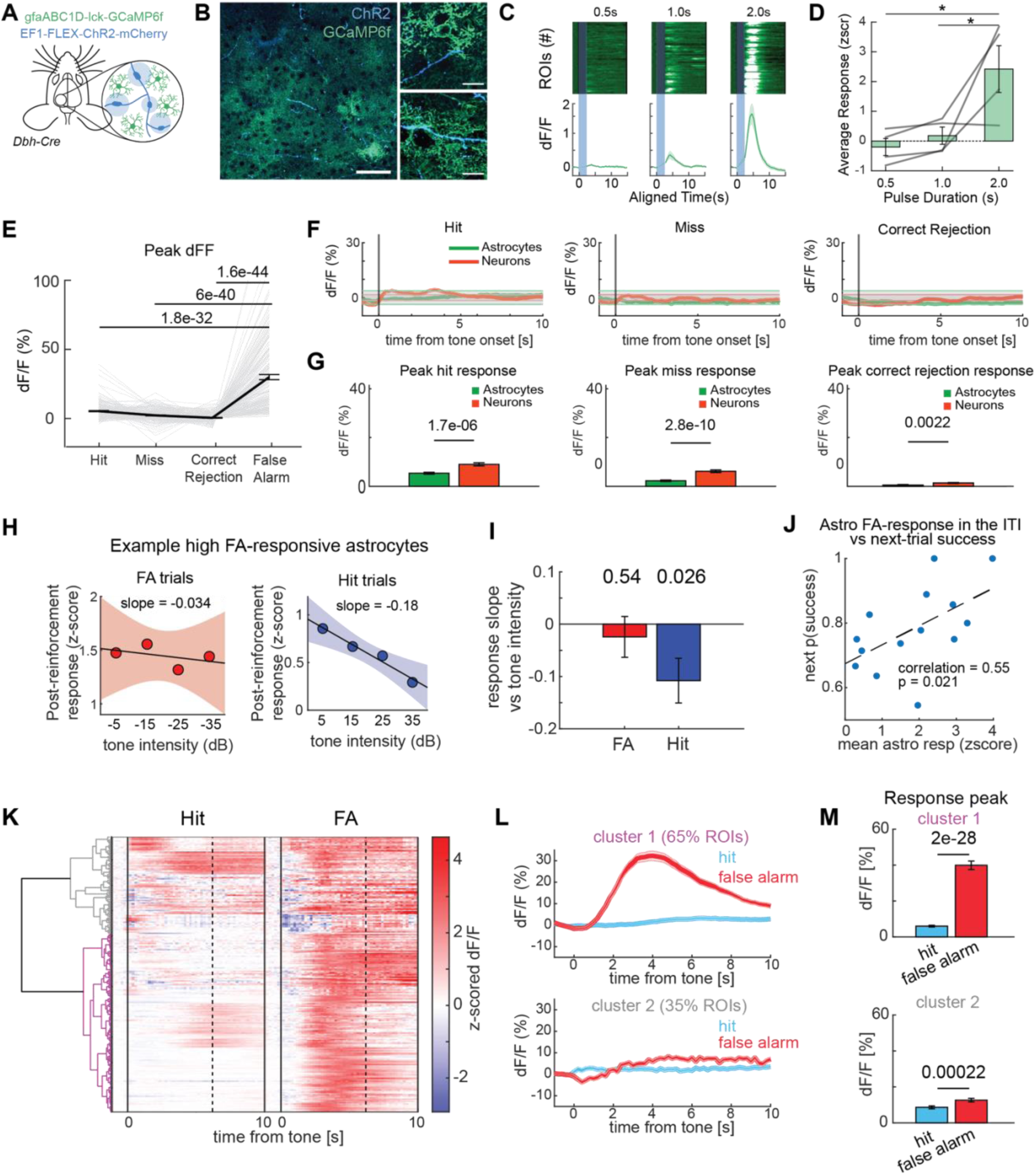
Astrocyte responses to LC activation and response dynamics in different trial types, relationship to next-trial behavior, and clustering analysis of astrocyte calcium dynamics in false alarm trials. **(A)** Schematic of experimental design for LC axon stimulation and astrocyte calcium imaging. **(B)** Example two-photon images of LC axons expressing ChR2 (blue) and astrocytes expressing GCaMP6f (green). Scale bar (left panel) = 50 µm; scale bar (right panels) = 20 µm. **(C)** Example astrocyte calcium traces following NE axon stimulation for 0.5, 1.0, and 2.0 seconds. **(D)** Average astrocyte calcium responses following NE axon stimulation for 0.5, 1.0, and 2.0 seconds. Each gray line indicates data from one animal. (n = 4 mice). **(E)** Peak dF/F for astrocytes during hit, miss, correct rejection, and false alarm trials in go/no-go task. Each gray line indicates one astrocyte. Black line indicates population average. Error bars are SEM. **(F)** Astrocyte and neuronal population dF/F on hit, miss, and correct rejection trials. n = 218 astrocytes, n = 4 mice; n = 208 neurons, n = 3 mice. **(G)** Average peak dF/F for hit, miss, and correct rejection trials for astrocytes and neurons. **(H)** Z-scored post-reinforcement calcium response of two example high FA-responsive astrocytes (top 50%, median split of peak FA responses) as a function of no-go tone intensity for false alarm (left) and go tone intensity for hit (right) trials. Dots indicate mean response for each tone intensity. Solid lines show least-squares linear regression fit; Shaded areas indicate the 95% confidence interval. **(I)** Slope of the z-scored astrocytes response vs no-go (FA) or go (hit) tone intensity for high FA-responsive astrocytes. n = 109 astrocytes, n = 4 mice. **(J)** Correlation between z-scored mean astrocytes response after a false alarm and next trial behavioral performance. Mean astrocyte activity is computed [-1,0] s from minimum next tone onset. Each dot is a session (n = 14), dashed line shows the best linear fit. **(K)** Z-scored, baseline subtracted, astrocyte responses to hits and false alarms sorted by hierarchical clustering. Clustering dendrogram shown on the left; branches corresponding to two identified clusters are shown in purple and grey. Black vertical dashed lines show the median time of next tone onset. n = 218 astrocytes, n = 4 mice. **(L)** Average response for hit and false alarm trials within each identified cluster. **(M)** Comparison of hit and false alarm peak astrocyte response within each cluster. P values show comparisons based on Mann-Whitney U-test in **D** (*, p<0.05), two-tailed paired t-test with Bonferroni correction in **E**, two-tailed unpaired t-test in **G**, two-tailed one sample t-test in **I**, and two-tailed paired t-test in **M.** Data show mean ± SEM.

**Figure S3.**
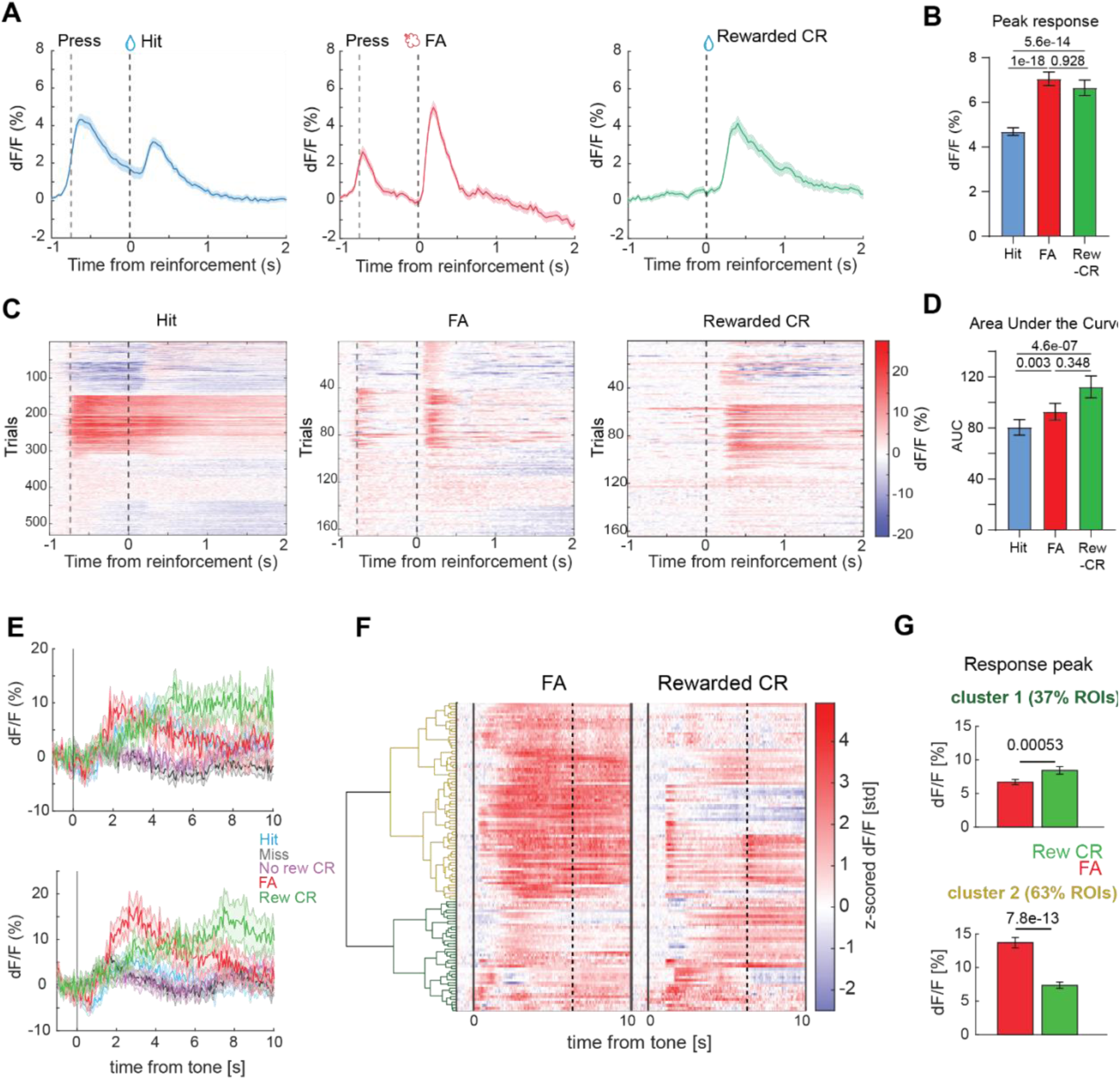
Cortical NE release and astrocyte calcium dynamics in response to a surprising reward in correct rejection trials. **(A)** Mean GRAB_NE2h_ dF/F signal on hit, false alarm, and rewarded correct rejection trials, aligned to the time of reinforcement. Solid lines and shaded areas display mean ± SEM. Dashed lines denote the time of lever push and reinforcement. **(B)** Post-reinforcement peak amplitude of GRAB_NE2h_ responses is higher after rewarded correct rejection and false alarm compared to hit trials. **(C)** Mean GRAB_NE2h_ dF/F responses across individual hit, false alarm and, rewarded correct rejection trials aligned to the time of reinforcement. **(D)** The AUC of GRAB_NE2h_ responses after reinforcement of rewarded correct rejection compared to hit and false alarm trials. Hits: n = 531 trials (n = 3 mice); FA: n = 165 trials (n = 3 mice); rewarded CR: n = 164 trials (n = 3 mice). **(E)** Example traces for hit, miss, correct rejection, and false alarm trials from two astrocytes. **(F)** Z-scored, baseline subtracted, average astrocyte dF/F for reinforced trial outcomes sorted by hierarchical clustering. Clustering dendrogram shown on the left; branches corresponding to two identified clusters are shown in green and yellow. **(G)** Comparison of false alarm and rewarded CR peak astrocyte response within each cluster. n = 124 astrocytes (n = 4 mice). P values show comparisons based on one-way ANOVA with Bonferroni’s post-hoc test in **B**, **D,** and two-tailed paired t-test in **G**. Data show mean ± SEM.

**Figure S4.**
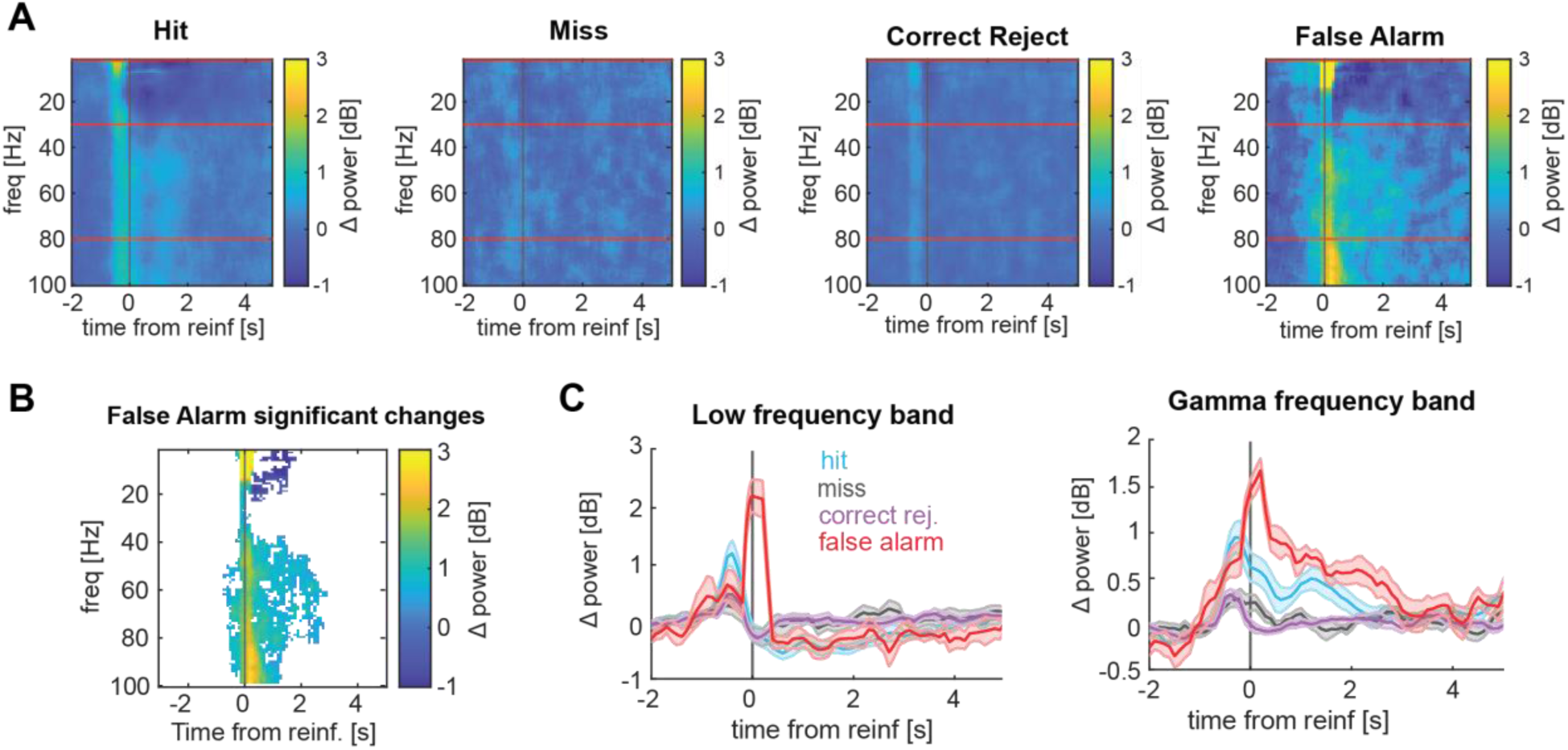
LFP power spectral analyses. **(A)** Time-frequency representations (TFR) of LFP power change relative to baseline, aligned to reinforcement delivery, for specific trial types (from left to right: hit, miss, correct rejection, and false alarm). Horizontal white dashed lines indicate gamma band frequency range limits. Pre-reinforcement power increases reflect tone responses and the 0.5 s moving window used for TFR computation. **(B)** Significant LFP power changes in false alarm trials derived using cluster-based permutation statistics (p<0.05). **(C)** Average low (2-30 Hz, left) and gamma (30-80 Hz, right) frequency band LFP power change from baseline for specific trial types.

**Figure S5.**
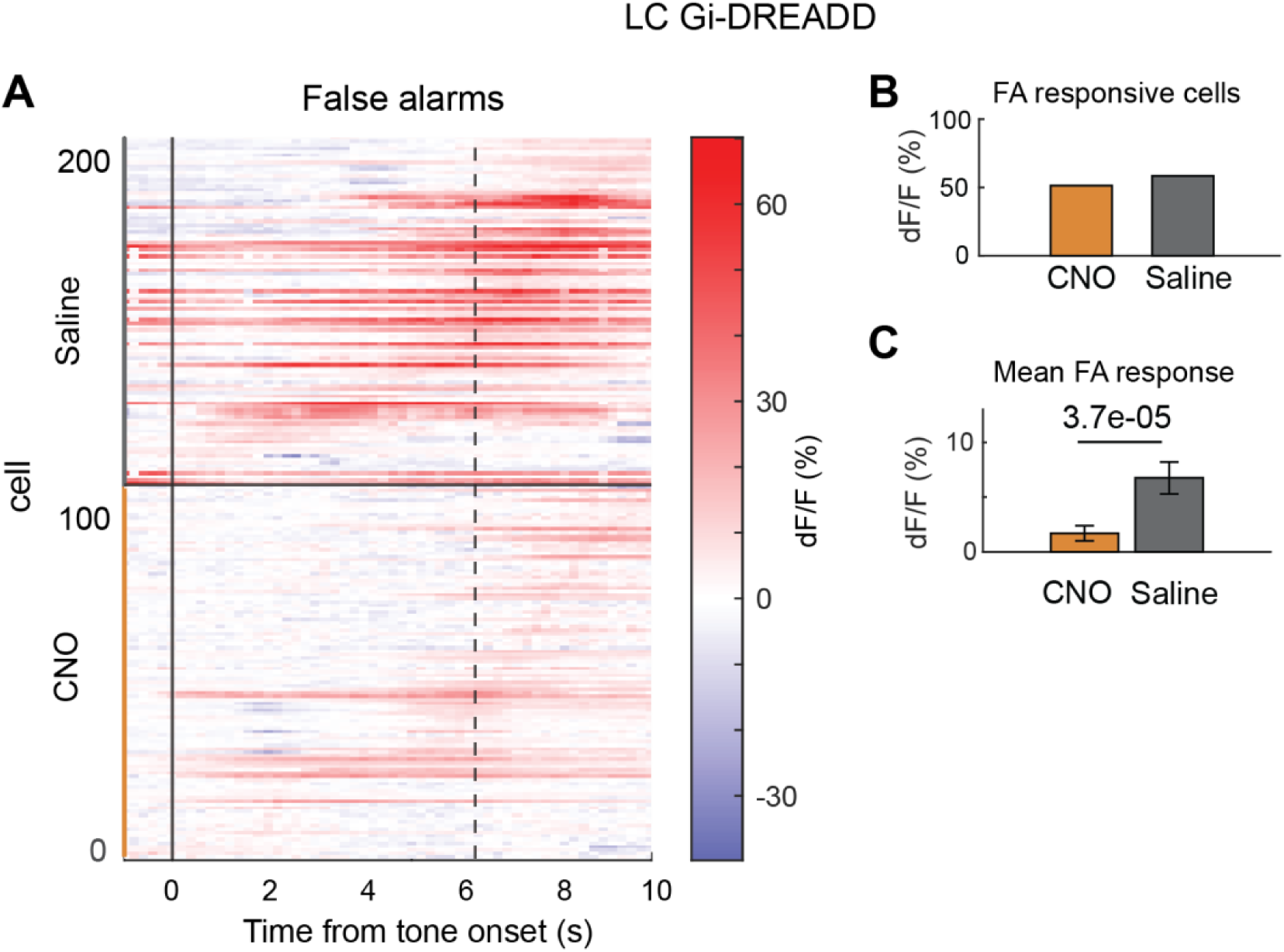
Chemogenetic silencing of LC neurons and astrocyte calcium dynamics. **(A)** Average astrocyte dF/F on false alarm trials for chemogenetic silencing of LC-NE in saline (top) and CNO (bottom) sessions. n = 101 astrocytes, saline; n = 108 astrocytes, CNO (n = 3 mice). Black vertical dashed lines show the median time of next tone onset**. (B)** Fraction of astrocytes responsive to false alarms in saline and CNO sessions. **(C)** Average mean dF/F for astrocytes in saline and CNO sessions. P values show comparisons based on two-tailed unpaired t-test. Data show mean ± SEM.

**Figure S6.**
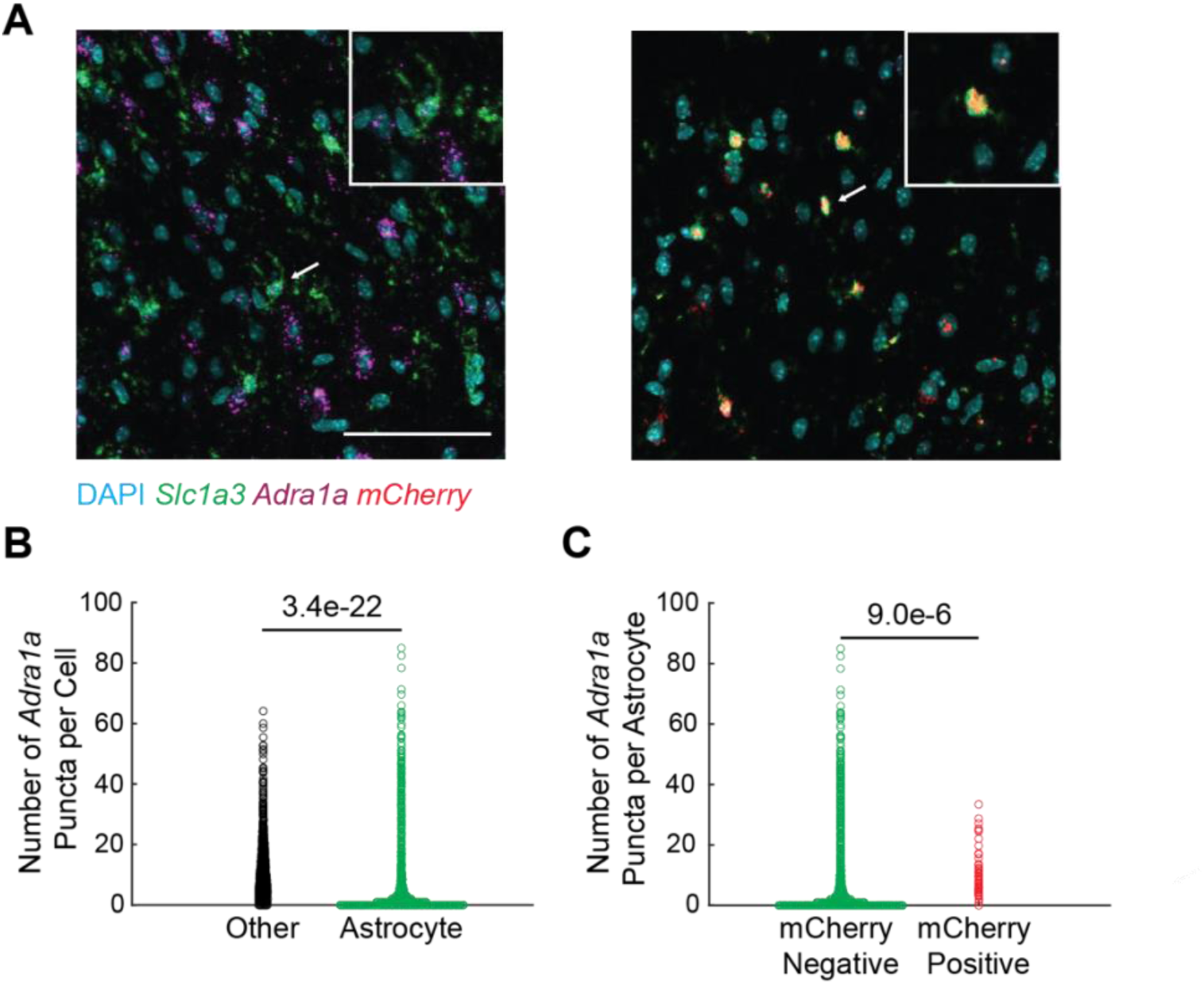
Adra1a knockdown in the cortex. **(A)** Example FISH images of the PFC of an Adra1a fl/fl mouse injected with AAV5-GFAP(0.7)-mCherry-T2A-iCre. (Left) A region outside of the injection site. The white arrow indicates a *mCherry* negative astrocyte (*Slc1a3* positive) highlighted in the inset panel. (Right) A region within the injection site. The white arrow indicates a *mCherry* positive astrocyte, putatively expressing *Cre*, highlighted in the inset. Scale bar = 100 µm. **(B)** Quantification of *Adra1a* puncta expression across mCherry negative astrocytes (*Slc1a3* positive) and non-astrocytic cells (n = 3 mice, n = 11,376 other cell types and 7,261 astrocytes). **(C)** Quantification of *Adra1a* puncta expression across *mCherry* positive and negative astrocytes (n = 3 mice, n = 7,261 *mCherry* negative astrocytes (same data as in **B**) and 593 *mCherry* positive astrocytes). P values show comparisons based on two-tailed unpaired t-test.

**Figure S7.**
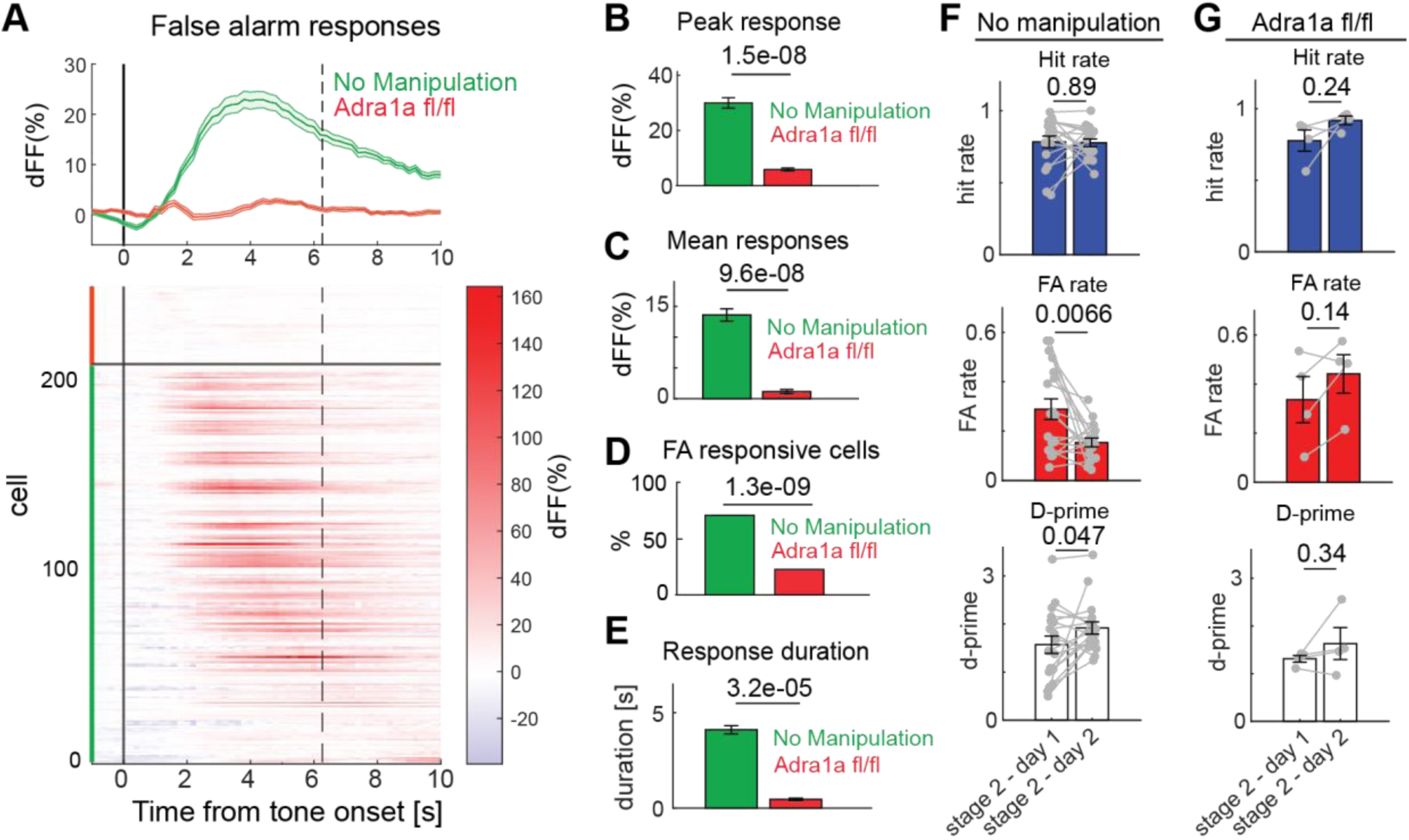
Astrocyte calcium dynamics in Adra1a KD mice. **(A)** Average astrocyte dF/F on false alarm trials without manipulations (green) and with knockdown of Adra1a receptors (red) sessions. The “no manipulation” mice are wild-type mice injected with gfaABC1D-GCaMP6f and Syn-jRGECO1a in the motor cortex. Population average (top) and individual cells (bottom); n = 218 astrocytes with no manipulation (n = 4 mice); n = 44 astrocytes with knock down of Adra1a (n = 3 mice). Black vertical dashed lines show the median time of next tone onset. **(B)** Average mean dF/F for astrocytes in no manipulation and knockdown of Adra1a sessions. **(C)** Average peak dF/F response in the two conditions. **(D)** Fraction of astrocytes responsive to false alarms in no manipulation and knockdown of Adra1a sessions. **(E)** Average false alarm response duration in the two conditions. **(F)** Hit rate (top), false alarm rate (middle), and d-prime (bottom) comparisons between the first and second day of training in the second stage of the task for animals with no manipulation (n = 18; stage two analyses limited to the first two days, the minimum duration across animals). **(G)** Same as **F** for Adra1a knockdown mice (n=4). P values show comparisons based on two-tailed unpaired t-test in **B, C, D**, two-tailed normal approximation to binomial test in **E,** and two-tailed paired t-test in **F, G**. Data show mean ± SEM.

**Figure S8.**
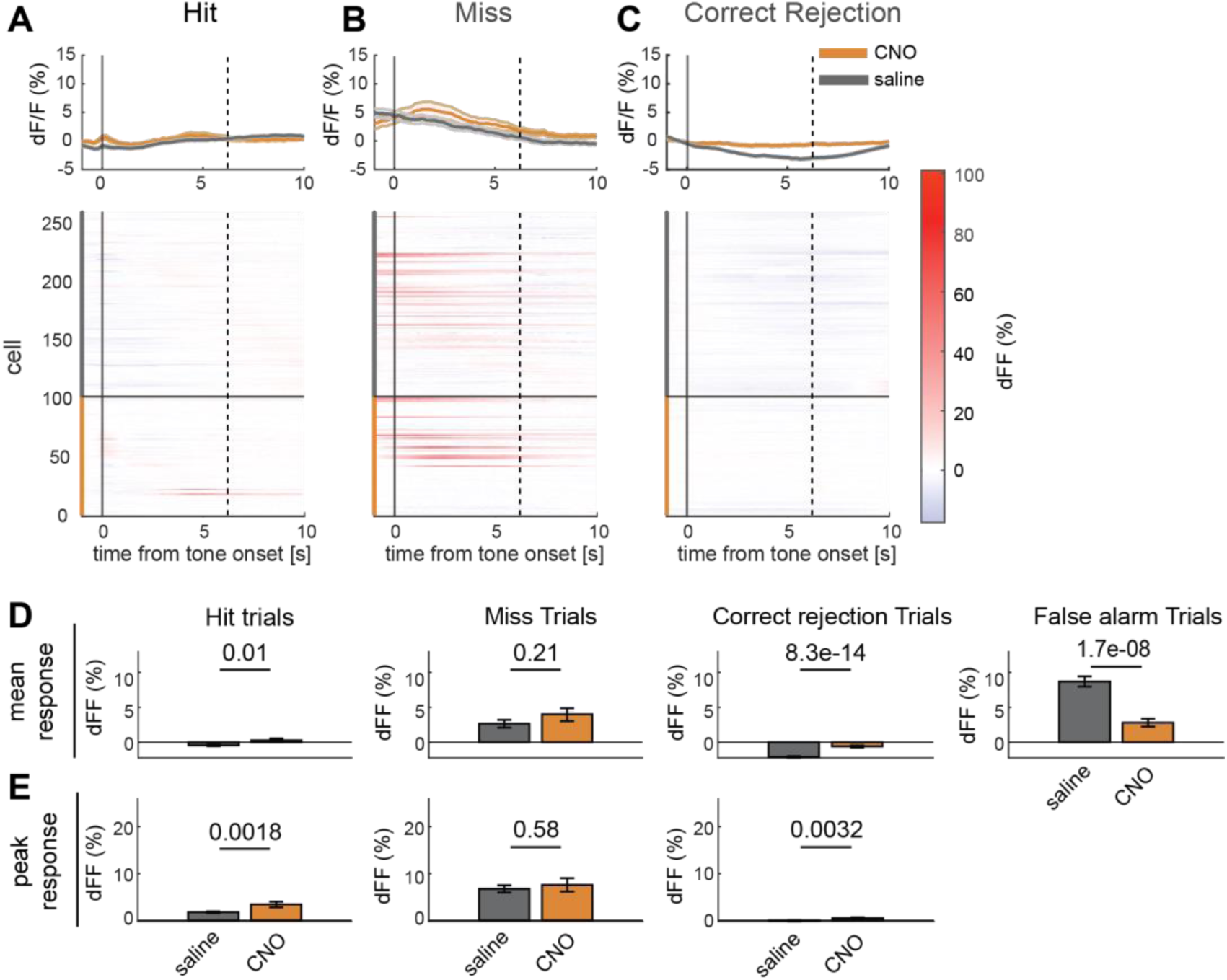
Chemogenetic manipulation of astrocyte calcium dynamics and astrocyte response profiles to different trial types. **(A)** Average astrocyte dF/F on hit trials for chemogenetic inactivation of astrocytes in saline (gray) and CNO (orange) sessions; population average (top) and individual cells (bottom). n = 160 astrocytes, saline; n = 99 astrocytes, CNO (n = 5 mice). **(B)** Same as **A** on miss trials. **(C)** Same as **A** on correct rejection trials. **(D)** Average mean dF/F for astrocytes in saline and CNO sessions for, from left to right: hit, miss, correct rejection, and false alarm trials. **(E)** Average peak dF/F for astrocytes in saline and CNO sessions, from left to right: hit, miss, and correct rejection trials. P values show comparisons based on two-tailed unpaired t-test in **D** and **E**. Data show mean ± SEM. Black vertical dashed lines in **A**, **B**, **C** show the median time of next tone onset.

**Figure S9.**
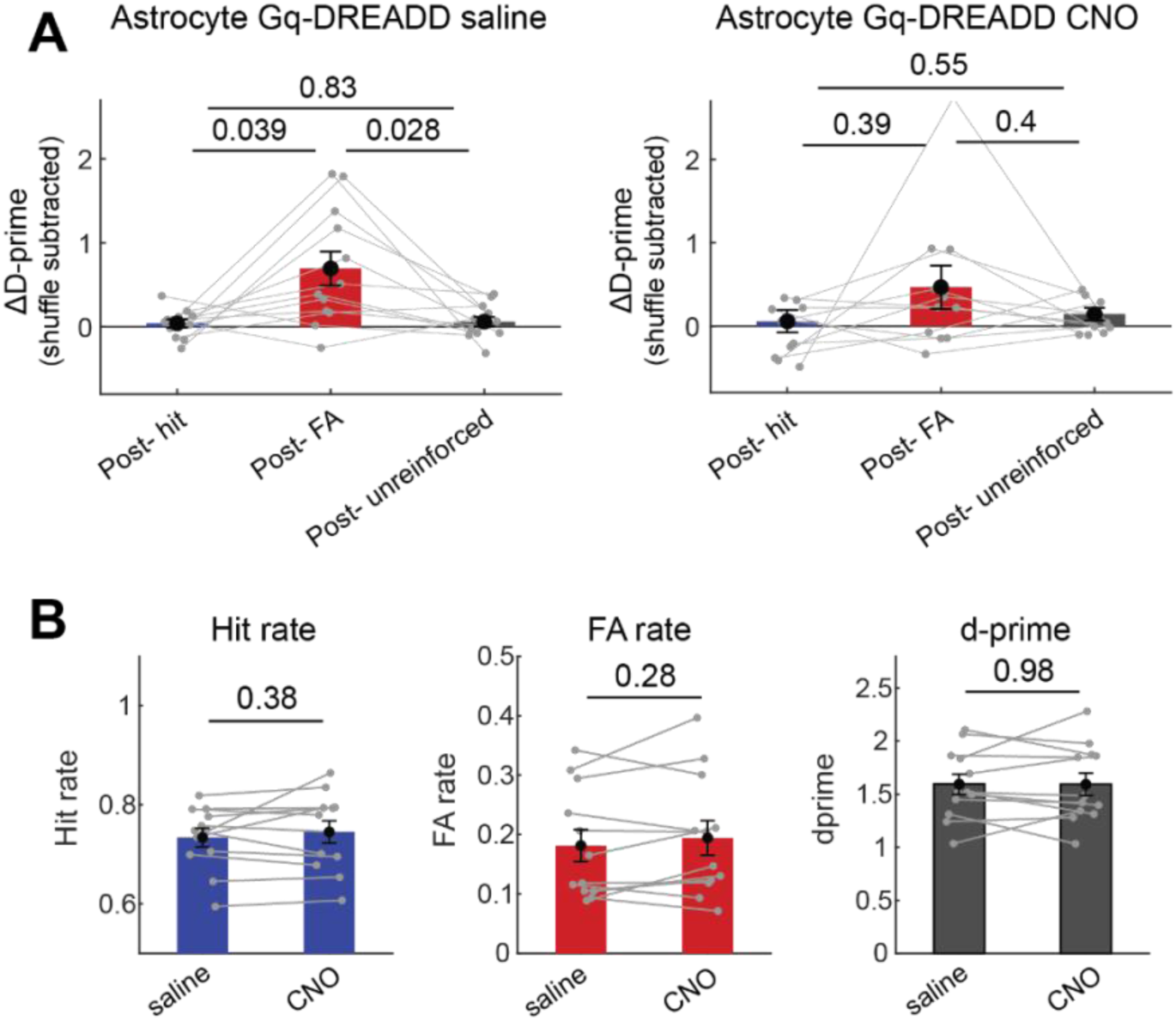
Behavioral metrics of Astro Gq-DREADD mice. **(A)** Change in d-prime after hit, false alarm, and unreinforced trials for saline (left) and CNO (right) sessions in astrocyte Gq-DREADD expressing animals. **(B)** Overall hit rate (left), false alarm rate (middle), and d-prime (right) in saline and CNO sessions. P values show comparisons based on two-tailed paired t-test. Data show mean ± SEM across n = 12 mice.

**Figure S10.**
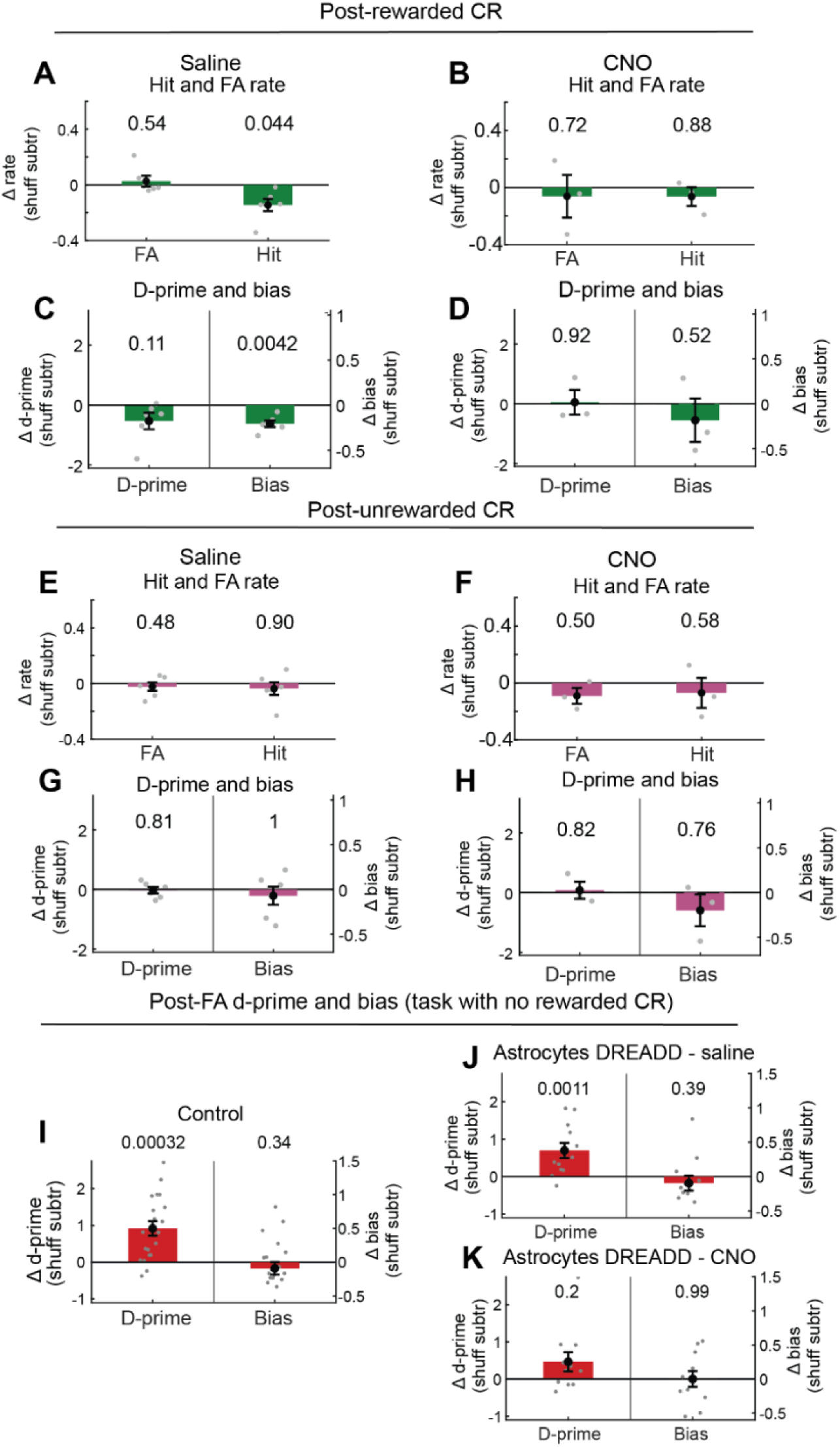
Behavioral effects after a rewarded correct rejection trial versus other trial types in control and chemogenetic astrocyte manipulation sessions. **(A)** Change in false alarm rate and hit rate on trials following a rewarded correct rejection for saline sessions, calculated after subtracting shuffled data. **(B)** Same as **A** for CNO sessions. **(C)** Change in d-prime (discriminability) and bias (tendency to press regardless of evidence) on trials following a rewarded correct rejection for saline sessions, calculated after subtracting shuffled data. **(D)** Same as **C** for CNO sessions. **E-H**, Same as **A-D** on trials following unrewarded CR. n = 6 mice for saline sessions; n = 3 mice for CNO sessions. **(I)** Change in d-prime and bias on trials following a false alarm for control animals in the task with no rewarded correct rejection (n = 20 mice). (**J, K**) Same as **i** for Astrocyte DREADD mice with saline (**J**) or CNO (**K**) injections. (n = 12 mice). P values show comparisons based on two-tailed one-sample t-test. Data show mean ± SEM.

**Figure S11.**
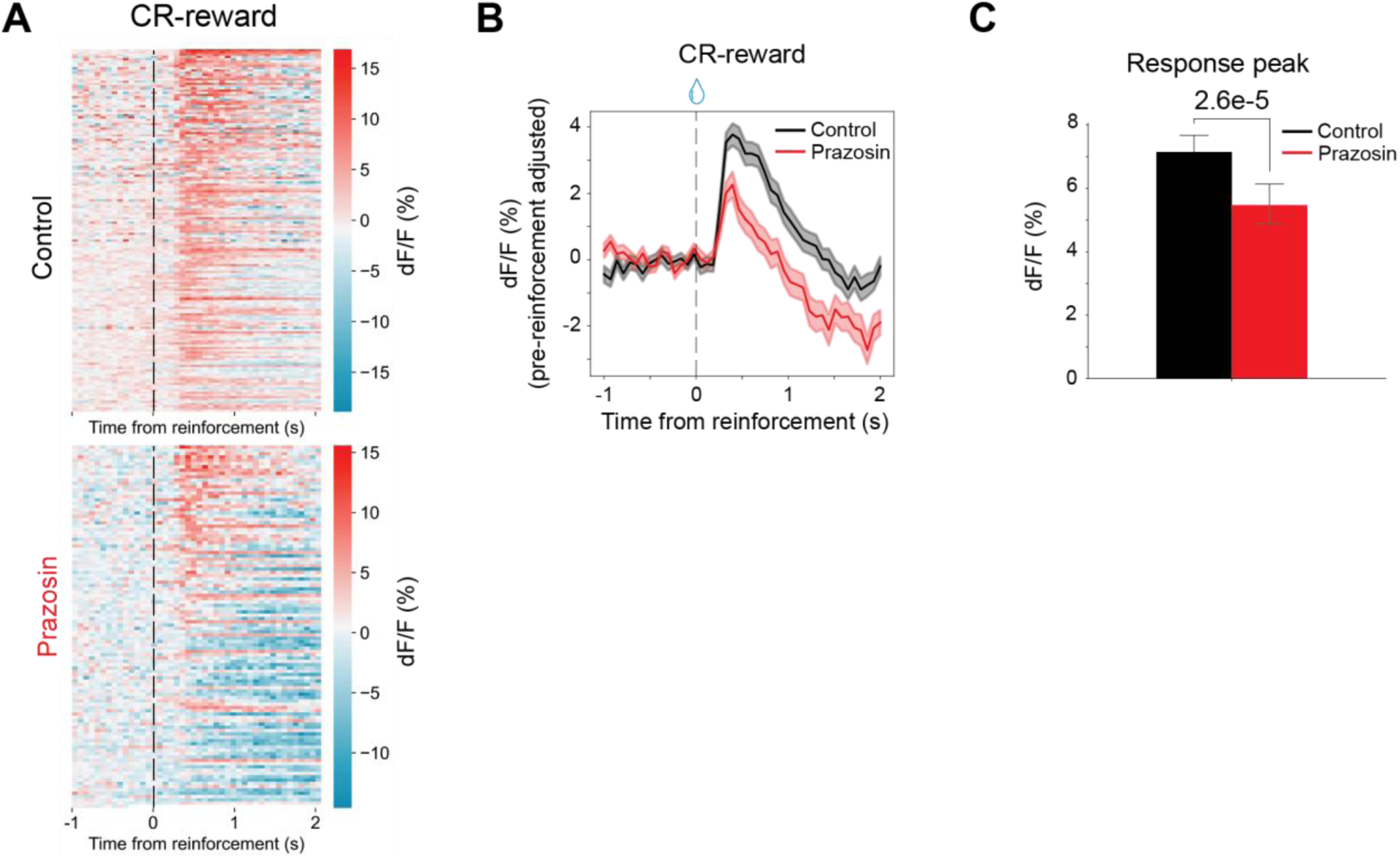
ATP dynamics in rewarded correct rejection trials in control and prazosin conditions. **(A)** Heatmaps of averaged GRAB_ATP_ signals aligned to time of reinforcement in control (top) and prazosin (bottom) conditions. **(B)** Average dF/F of control (black) and prazosin (red) conditions. **(C)** Magnitude of post-reinforcement response peaks. Control: n = 162 trials from 3 mice. Prazosin = 110 trials from 3 mice. P value was computed using two-tailed unpaired t-test in **C**. Data show mean ± SEM.

**Figure S12.**
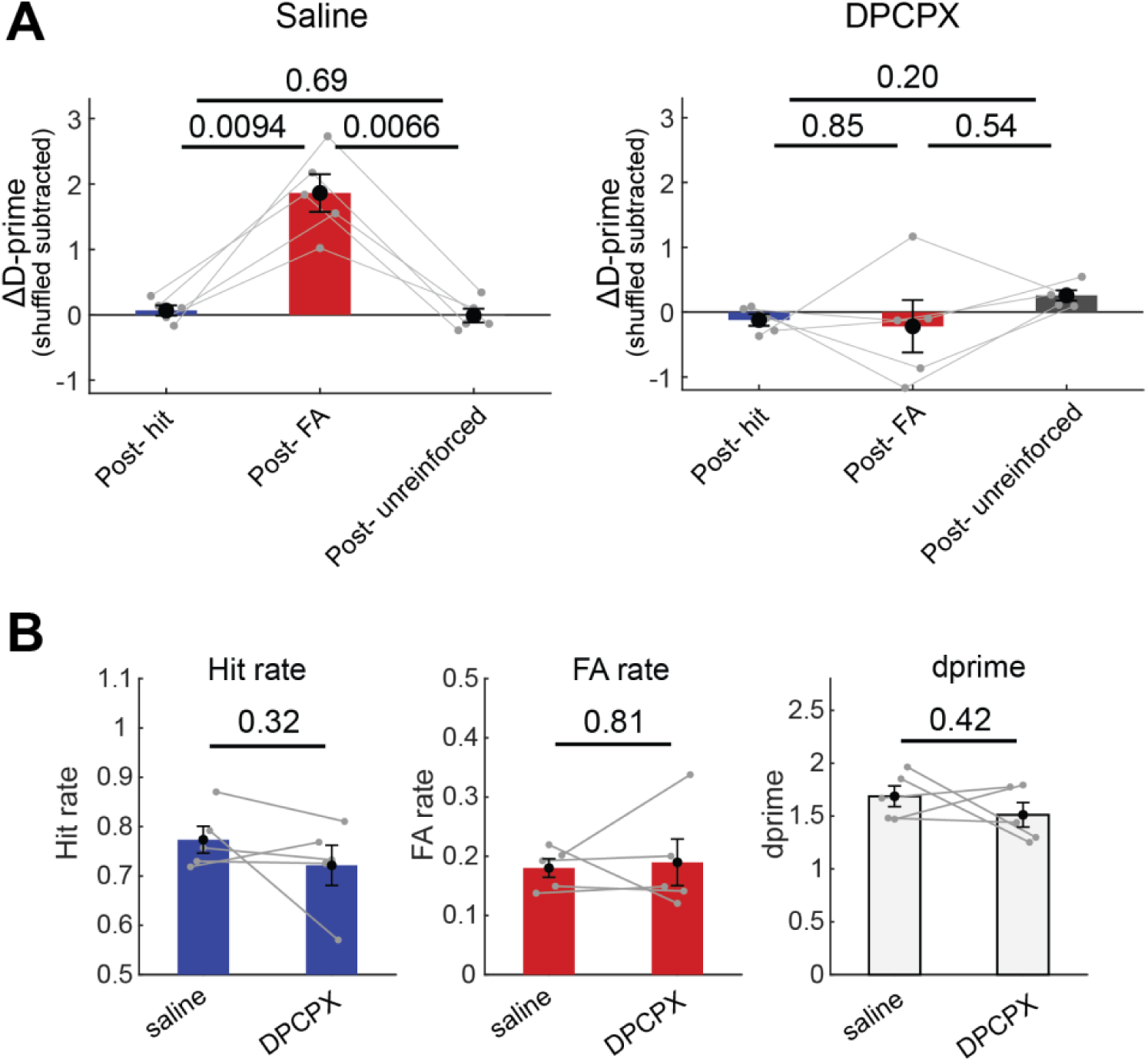
Behavioral metrics of mice with DPCPX application. **(A)** Change in d-prime after hit, false alarm, and unreinforced trials for saline (left) and DPCPX (right) sessions. **(B)** Overall hit rate (left), false alarm rate (middle), and d-prime (right) in saline and DPCPX sessions. P values show comparisons based on two-tailed paired t-test. Data show mean ± SEM across n = 5 mice.

**Figure S13.**
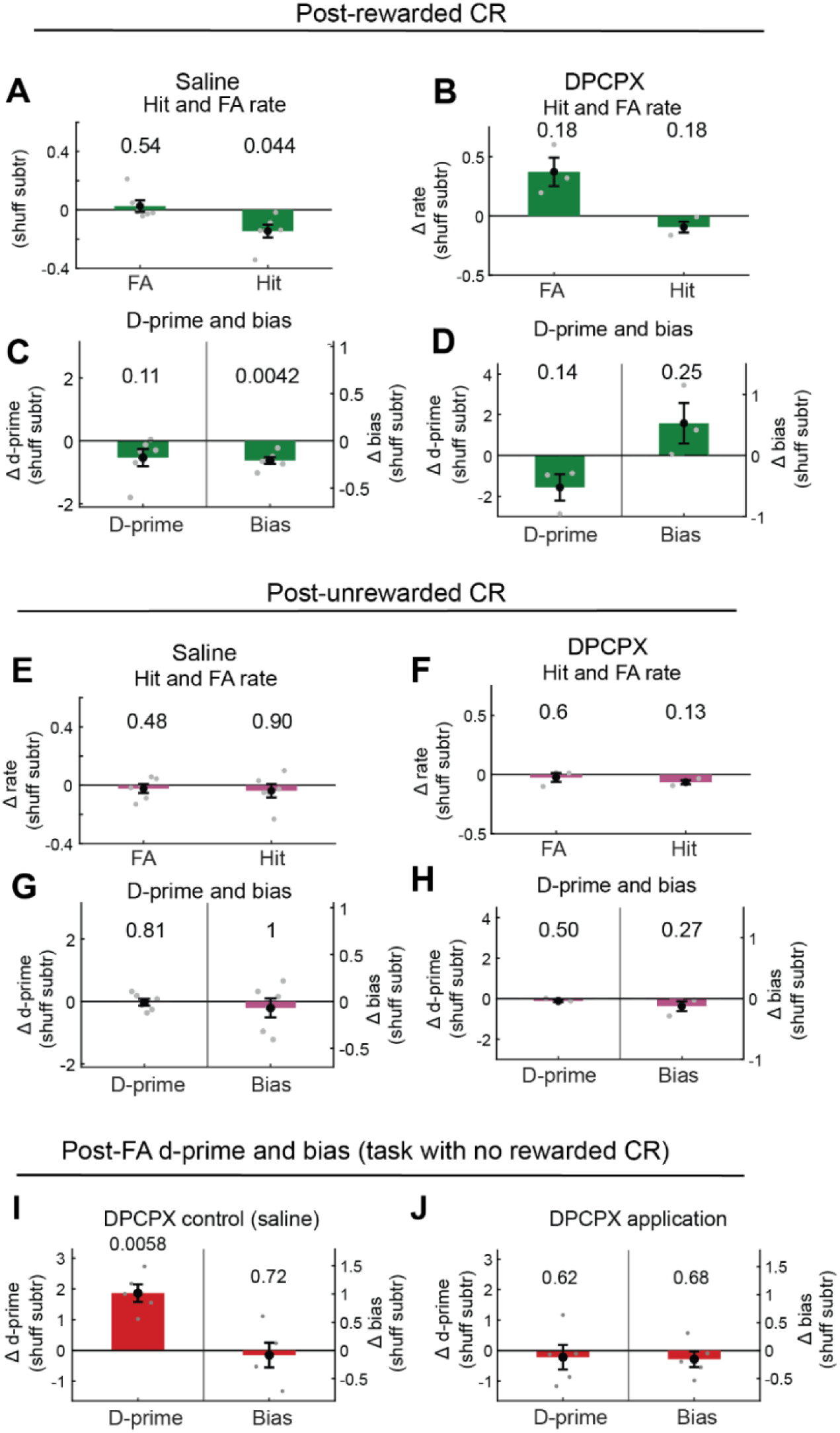
Behavioral effects after a rewarded correct rejection trial versus other trial types in control and DPCPX sessions. **(A)** Change in false alarm rate and hit rate on trials following a rewarded correct rejection for saline sessions, calculated after subtracting shuffled data. **(B)** Same as **A** for DPCPX sessions. **(C)** Change in d-prime (discriminability) and bias (tendency to press regardless of evidence) on trials following a rewarded correct rejection for saline sessions, calculated after subtracting shuffled data. **(D)** Same as **c** for DPCPX sessions. (**E-H**) Same as **A-D** on trials following unrewarded CR. n = 6 mice for saline sessions; n = 3 mice for DPCPX sessions. (**I**, **J)** Change in d-prime and bias on trials following a false alarm for saline control (**I**) and DPCPX application (**J**) sessions in the task with no rewarded CR. (n = 5 mice). P values show comparisons based on two-tailed one-sample t-test. Data show mean ± SEM.

**Figure S14.**
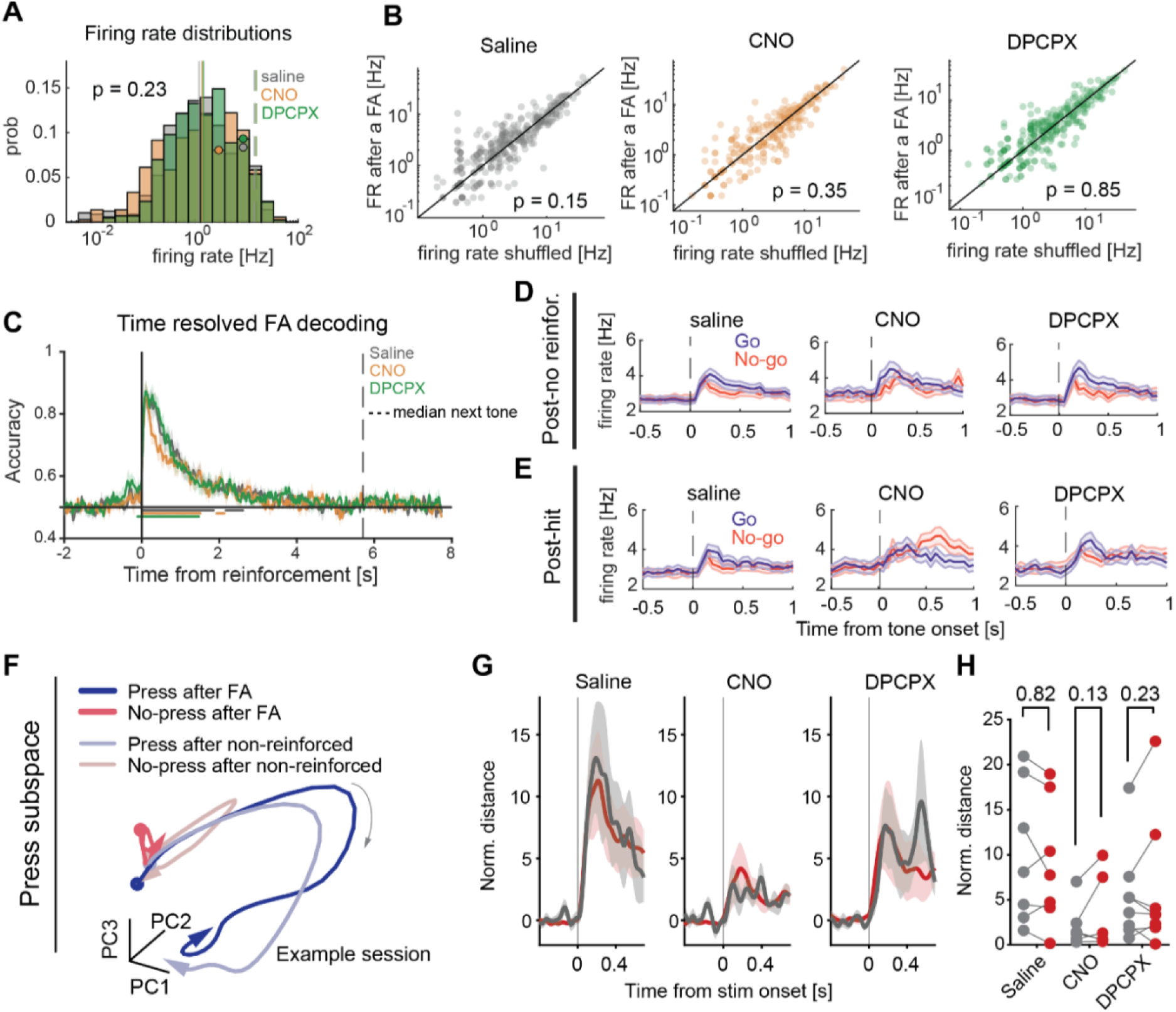
Neuronal activity and stimulus discriminability in post-hit, miss, and correct rejection trials and population encoding in post-false alarm trials across different conditions. **(A)** Firing rate distributions in the response window ([50,250] ms from tone onset) for saline (gray), CNO (orange), and DPCPX (green) sessions on a logarithmic scale. Solid lines show medians, dashed lines show 95^th^ percentiles. Dots indicate the bins that example neurons shown in Figure 4A belong to. **(B)** Scatter plots of mean firing rate in the response window after a false alarm response compared to shuffled data for saline (left), CNO (middle), and DPCPX (right) sessions. Data are plotted on logarithmic axes. **(C)** Time-resolved decoding accuracy of false alarm trials from neuronal population activity (25 neurons randomly selected 50 times) across saline, CNO, and DPCPX sessions. **(D)** Average neuronal firing rate for go (blue) and no-go (red) stimuli following unreinforced trials in saline (left), CNO (middle), and DPCPX (right) sessions. **(E)** Same as **D** following hit trials. n = 526 neurons (n = 6 mice); n = 360 neurons, CNO (n = 5 mice); n = 429 neurons, DPCPX (n = 5 mice). **(F)** Example neuronal population trajectories projected into the first three PCs of press-related subspace. Each color corresponds to a distinct trial type. **(G)** Time-resolved normalized Euclidean distance between press and no-press trials in saline, CNO, and DPCPX sessions. **(H)** Comparison of normalized Euclidean distance between post-false alarm (red) and post-unreinforced (gray) trials in saline, CNO, and DPCPX sessions. n = 7 sessions, saline; n = 5 sessions, CNO; n = 9 sessions, DPCPX. P values show comparisons based on two-tailed Kruskal-Wallis test in **A**, on two tailed Wilcoxon signed rank test in **B**, and on one-tailed paired t-test in **H**. Data show mean ± SEM.

**Figure S15.**
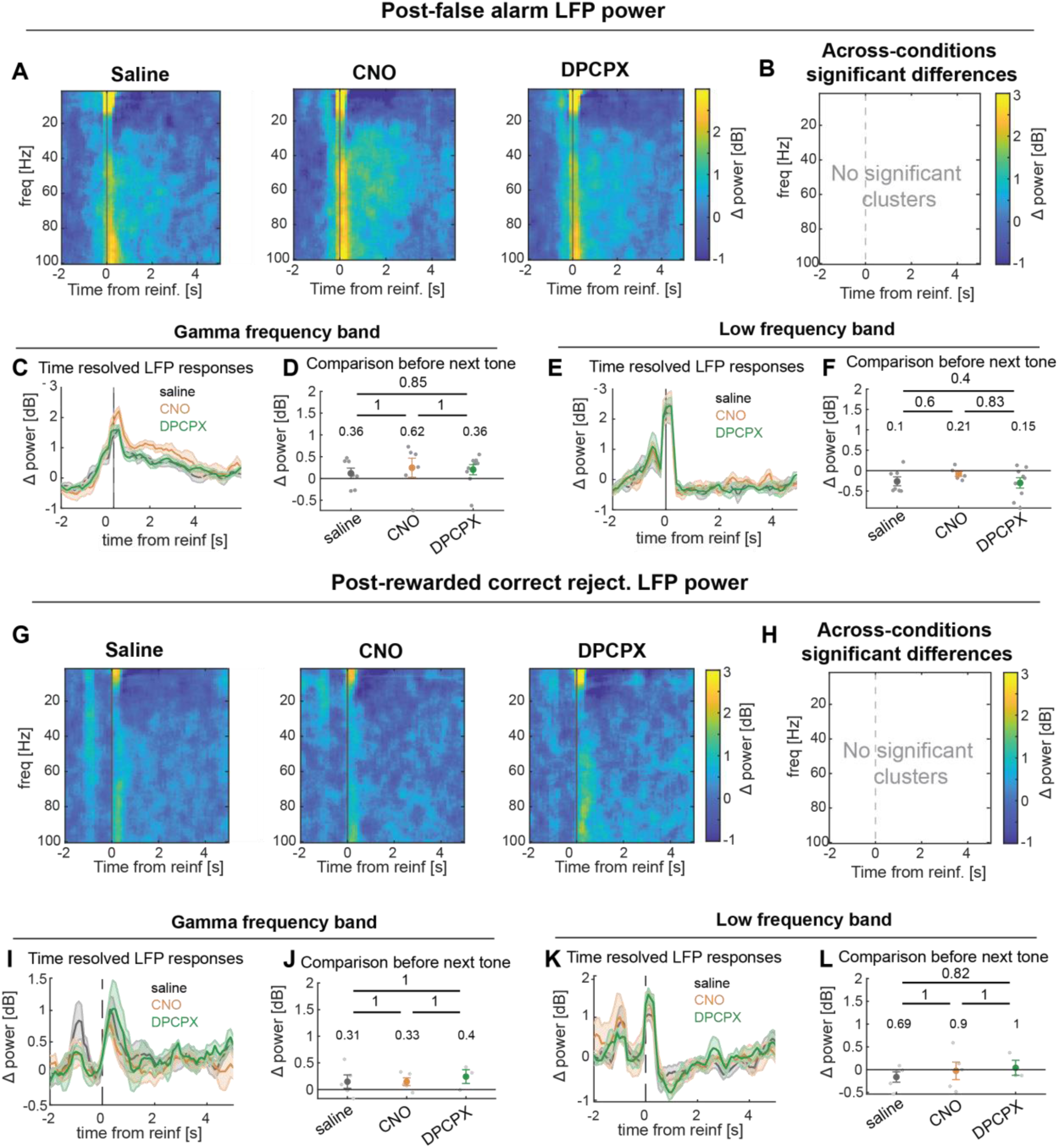
LFP power spectral comparison across different conditions. **(A)** Time-frequency representations of LFP power change relative to baseline on false alarm trials, aligned to reinforcement delivery, for saline (n = 7), CNO (n = 5), and DPCPX (n = 9) sessions. **(B)** Significant differences across conditions, assessed with a cluster-based permutation test on one-way ANOVA (session labels shuffled, p<0.05). **(C)** Average gamma (30-80Hz) frequency band LFP power change from baseline on false-alarm trials for saline, CNO, and DPCPX sessions. **(D)** Comparison across gamma frequency band LFP power changes [-1,0] s from minimum next tone onset vs baseline and across conditions. (**E, F)**, same as **C, D** for low frequency band. **(G-L)** same as **A-F** for correct rejection-reward trials. P values in **D, F, J, L** show comparisons based on two-tailed one-sample t-test (vs baseline) or unpaired t-test (across conditions). Data show mean ± SEM.

**Figure S16.**
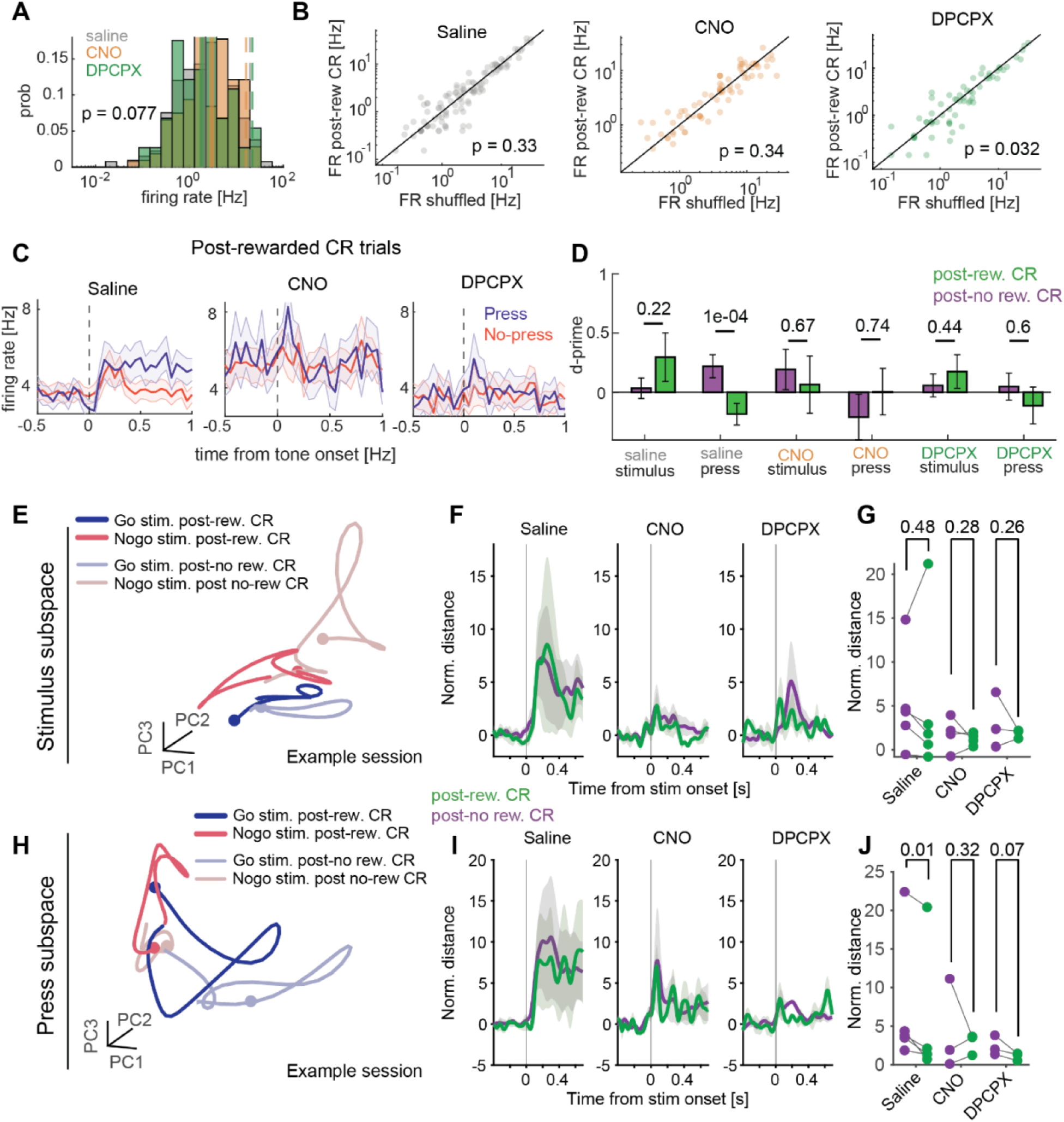
Neuronal activity, press discriminability, and population encoding in post-rewarded correct rejection trials across different conditions. **(A)** Firing rate distributions in the response window ([50,250] ms from tone onset) for saline (gray), CNO (orange), and DPCPX (green) sessions on a logarithmic scale. Solid lines show medians, dashed lines show 95^th^ percentiles. **(B)** Scatter plots of mean firing rate in the response window after a false alarm compared to shuffled data for saline (left), CNO (middle), and DPCPX (right) sessions. Data are plotted on logarithmic axes. **(C)** Average neuronal firing rate for press (blue) and no-press (red) trials following rewarded correct rejection trials in saline (left), CNO (middle), and DPCPX (right) sessions. **(D)** Comparison of stimulus and press neuronal d-prime between trials following a rewarded (green) and an unrewarded (purple) correct rejection for saline, CNO, and DPCPX sessions. n = 133 neurons (n = 4 mice); n = 52 neurons, CNO (n = 3 mice); n = 74 neurons, DPCPX (n = 3 mice). **(E)** Example neuronal population trajectories projected into the first three PCs of stimulus-related subspace. Each color corresponds to a distinct trial type. **(F)** Time-resolved normalized Euclidean distance between go- and no go-stimulus trials in saline, CNO, and DPCPX sessions. **(G)** Comparison of normalized Euclidean distance between post-rewarded correct rejection (green) and post-unrewarded correct rejection (purple) trials in saline, CNO, and DPCPX sessions. **(H-J)** Same as **E-G**, but for press-related population activity. n = 5 sessions, saline; n = 4 sessions, CNO; n = 3 sessions, DPCPX. P values show comparisons based on two-tailed Kruskal-Wallis test in **A** Wilcoxon signed rank test in **B**, two-tailed paired t-test in **D**, and one-tailed paired t-test in **G, J**. Data show mean ± SEM.

**Table S1.** Summary of across-session Pearson correlations between each signal (reinforcement-aligned responses, averaged [-1,0] s relative to the minimum next tone onset) and the probability of a successful response on the next trial. Significant correlations are highlighted with bold font. Analyses include 7-14 sessions across conditions.

| Brain signal | Pearson Correlation | P-value |
| --- | --- | --- |
| Mean astrocyte response – FA trials | **0.55 ± 0.21** | **0.021** |
| Mean astrocyte response – no-reinf. trials | -0.11 ± 0.30 | 0.66 |
| Mean neuronal calcium response – FA trials | 0.26 ± 0.54 | 0.31 |
| Gamma power – FA trials | -0.07 ± 0.50 | 0.56 |
| Low power – FA trials | -0.25± 0.47 | 0.30 |
| Pupil diameter change – FA trials | 0.27± 0.23 | 0.14 |
| Neural population FA signals (NPX decoding) | -0.05 ± 0.58 | 0.54 |

